# Data-driven strategies for drug repurposing: insights, recommendations, and case studies

**DOI:** 10.1101/2025.08.18.670836

**Authors:** Susanna Savander, Nurettin Nusret Curabaz, Amna Mumtaz Abbasi, Asifullah Khan, Khalid Saeed, Ziaurrehman Tanoli

## Abstract

Drug discovery is a complex, time-intensive, and costly process, often requiring more than a decade and substantial financial investment to bring a single therapeutic to market. Drug repurposing, the systematic identification of new indications for existing approved drugs, offers a cost-effective and expedited alternative to traditional pipelines, with the potential to address unmet clinical needs. In this study, we present a comparative analysis of drug–target interaction data from three extensively curated resources: ChEMBL, BindingDB, and GtoPdb, evaluating their release histories, curation methodologies, and coverage of approved and investigational compounds and targets. To facilitate therapeutic interpretation, we manually classified ChEMBL targets into 12 high-level biological families and mapped 817 clinically approved drug indications into 28 broader therapeutic groups. This structured framework enabled a systematic profiling of physicochemical properties among approved drugs across therapeutic categories. Our analyses revealed associations between physicochemical characteristics and therapeutic groups, providing practical guidance for indication-specific compound prioritization and refining the repurposing studies. We also examined cross-indication drug approvals to identify areas with high repurposing potential. Finally, we implemented a pathway-based computational pipeline to predict repositioning opportunities for FDA-approved drugs across ten major cancer types, demonstrating its adaptability to other disease contexts. Overall, this work consolidates drug-target data and computational repurposing into a data-driven framework that advances drug discovery and translational applications.

## 1. INTRODUCTION

The development of a new drug is an extraordinarily complex, resource-intensive, and time-consuming process. It typically requires $1–3 billion in investment and spans 9 to 15 years from target identification to market approval [1]. This multi-stage process begins with target identification, followed by hit identification, lead optimization, clinical evaluation (across three trial phases), and post-marketing surveillance (phase 4) presents numerous bottlenecks that collectively limit the pace of therapeutic innovations [2]. Recent advances in experimental screening technologies have partially mitigated these challenges. For instance, DNA-encoded libraries (DELs) enable ultra-high-throughput screening of millions of compounds against selected molecular targets [3]. DELs have revolutionized the hit identification stage by generating vast numbers of drug-target interaction (DTI) data points at minimal cost. However, the resulting data are largely qualitative (binary active/inactive) and provide limited insight into binding strength or selectivity. Complementary approaches such as Proteome Integral Solubility Alteration (PISA) assays extend this principle by assessing proteome-wide ligand-induced thermal stability shifts, thereby offering indirect quantitative information about binding affinity and target engagement [4]. Nevertheless, PISA remains experimentally demanding and low-throughput, making it unsuitable for large-scale compound screening.

These methodological contrasts, DELs providing breadth but limited depth, and PISA offering depth but limited scale, underscore a critical unmet need for computational frameworks that can integrate, extrapolate, and quantify DTIs efficiently. Computational approaches bridge the gap between experimental throughput and mechanistic resolution, enabling the prediction of binding affinities across large chemical and proteomic spaces. For example, MMAtt-DTA [5], an attention-based architecture, predicts binding affinities for over 452,000 compounds and 1,251 human protein targets with high accuracy. Similarly, TransDTI [6] evaluated a diverse array of protein embeddings, demonstrating the utility of Evolutionary Scale Modeling (ESM) [7] and AlphaFold-derived features [8] as effective alternatives to conventional BERT-style embeddings. Given their expanding scope, computational DTI prediction models must undergo rigorous experimental or benchmarking validation to ensure translational reliability. This principle is comprehensively discussed in Tanoli et al. [9], which outlines best practices and current standards for evaluating DTI model performance.

Most deep learning approaches in DTI prediction depend heavily on high-quality training data sourced from manually curated databases, including ChEMBL [10], BindingDB [11], IUPHAR/BPS Guide to Pharmacology (GtoPdb) [12], DrugTargetCommons [13], among others. As summarized in previous reviews [14, 15], dozens of publicly available DTI resources now exist, though their quality, coverage, and data curation standards vary significantly. In a recent landmark review published in Nature Reviews Drug Discovery [16], Tanoli et al. systematically categorized 181 in silico tools using a standardized ontology, classifying them into 15 pharmacologically relevant categories. Accompanying this work, a web-based catalogue, iDRC-R4A (https://www.idrc-r4a.com/index.php) is developed to provide up-to-date information on in silico tools. This catalogue includes expert-driven rankings (via 27 field specialists), in which ChEMBL, BindingDB, and GtoPdb are ranked among the high-quality DTI databases (with ranking scores of 10, 9, and 8, respectively). Collectively, these three resources are among the most mature and robust, with more than 20 public releases (**Supplementary Figure 1**) each over the past two decades. These databases provide comprehensive, expertly curated datasets that encompass both primary and off-target interactions. Given their depth, consistency, and expert curation, the present study focuses exclusively on ChEMBL, BindingDB, and GtoPdb, utilizing their datasets to enable a thorough exploration of DTI.

While efficacy is a critical parameter, lead optimization also requires a thorough evaluation of Absorption, Distribution, Metabolism, Excretion, and Toxicity (ADMET) properties [17]. In addition, physicochemical descriptors such as partition coefficient (alogP), hydrogen bond donors (HBD) and acceptors (HBA), as well as topological polar surface area, play a significant role in determining a compound’s viability in subsequent stages of drug development. Although this study does not explore ADMET characteristics, we analyzed physicochemical properties for indication-specific profiling of small molecules. This study also evaluates the drug repurposing potential of approved drugs across 28 high-level therapeutic indication groups. Drug repurposing has emerged as a cost-effective strategy to accelerate development timelines and address unmet medical needs, particularly in rare or neglected diseases [16]. Notable examples include minoxidil, originally an anti-hypertensive agent, now widely repurposed for male-pattern hair loss [18], and imatinib, first approved for chronic myeloid leukemia (CML) and later found effective against gastrointestinal stromal tumors (GISTs) through inhibition of the KIT (CD117) tyrosine kinase [19].

Expanding on this foundation, we integrated comparative DTI analyses from ChEMBL, BindingDB, and GtoPdb, and additionally annotated targets into 12 high-level families of targets and therapeutic indications into 28 groups to support systematic analysis. By examining the relationships between therapeutic indication groups and the physicochemical properties of their corresponding approved drugs, we identified distinct clustering patterns among indication groups and physicochemical properties that may guide the design of novel therapeutics tailored to specific indication groups. Additionally, through the analysis of overlapping approved drugs shared across multiple indication groups, we derived evidence-based recommendations for identifying new drug repurposing opportunities. Finally, we implemented a pathway-based case study to demonstrate drug repositioning potential across major cancer types, providing a reference computational framework that can be readily extended to other therapeutic indications in future research.

### Key points

1. Comparative curation strategies and drug–target interaction profiles across ChEMBL, BindingDB, and GtoPdb.
2. Systematic classification of targets (12 families) and drug indications (28 disease groups), providing a structured view of therapeutic space.
3. Indication-specific physicochemical property patterns and cross-indication drug overlaps that inform both novel drug design and evidence-based repurposing opportunities.
4. Data driven recommendations and case studies for drug repurposing.

## 2. MATERIALS AND METHODS

### 2.1. Curation strategies and general overview of ChEMBL, BindingDB and GtoPdb

ChEMBL, BindingDB, and the GtoPdb are three of the most comprehensive and widely utilized public repositories for DTI data with more than 25 releases each (**Supplementary Figure 1**). Collectively, they provide an essential foundation for computational pharmacology, drug repurposing, virtual screening, and the development of machine learning models for tasks such as binding affinity estimation, and compound prioritization.

ChEMBL, maintained by EMBL–EBI and first publicly released in 2009, has since progressed through 35 major version updates. It now contains over 21 million bioactivity measurements involving >2.4 million ligands and biological targets >16,000 targets [10]. A total of 7,110 compounds in ChEMBL v33 have max_phase > 0, comprising 3,492 approved drugs and 3,618 investigational compounds (small molecules: 6828, biological ligands: 260 and unknown:22, **Supplementary File 1**). **Supplementary Figure 2** presents the number of first approvals per decade for these small molecules and biologics (including oligosaccharides, proteins, and other modalities) recorded in the ChEMBL database. The highest number of small molecule approvals occurred in the 1980s (n = 314), while biologics peaked in the 2000s (n = 13).

BindingDB developed in the early 2000s at the University of North Carolina, focuses specifically on experimentally determined binding affinities (e.g., K_ⅈ_, Kd, IC_50_). The current dataset contains over 2.4 million measurements covering ∼1.3 million unique ligands and nearly 9 thousand targets [11]. GtoPdb, the successor to IUPHAR‐DB, became publicly accessible in 2014 and has since followed a quarterly release schedule. Its data are based on expert curation of pharmacological literature, emphasizing clinically relevant target classes such as GPCRs, ion channels, and nuclear receptors. As of the latest release, GtoPdb contains curated data on 3,039 targets and 12,163 ligands [12].

All three databases incorporate manual curation and accept direct data submissions from published literature. Chemical structures are represented using standardized formats such as SMILES, InChI, and InChIKey, with cross-references to external resources including PubChem [20], UniChem [21], and DrugBank [22]. Discrepancies in structure assignment, assay annotation, or target linkage are resolved through expert review by in-house teams. Version-controlled releases ensure data traceability, reproducibility, and transparency. **Supplementary Figure 1** shows the release timelines for ChEMBL, BindingDB, and GtoPdb. While release histories for ChEMBL and GtoPdb are directly obtained from their respective websites, BindingDB does not maintain publicly accessible version logs. To address this, we contacted the BindingDB team, who provided annual data growth statistics, enabling us to approximate its release trajectory. It is important to note that the analyses conducted in this study was initiated in 2024 and may not reflect the most recent updates to these databases.

To assess the journal coverage of drug–target activity data curated by ChEMBL, BindingDB, and GtoPdb, we extracted article-level metadata (PubMed IDs) from each database. In total, 80,286 articles are curated by ChEMBL (V33), 43,038 by BindingDB, and 8,183 by GtoPdb. Journal names corresponding to each PubMed ID are retrieved using the Entrez programming utilities (E-utilities) via the Biopython package. Among these, 2,360 articles are common to all three databases, while 38,912 are shared between ChEMBL and BindingDB, 104 between ChEMBL and GtoPdb, and 374 between GtoPdb and BindingDB (**Figure 1D**). ChEMBL curates articles from 215 unique journals, BindingDB from 197, and GtoPdb from 657. A comparative analysis of the top 10 most frequently cited journals in each database revealed considerable overlap; for example, Journal of Medicinal Chemistry, Bioorganic & Medicinal Chemistry Letters, Bioorganic & Medicinal Chemistry, and ACS Medicinal Chemistry Letters appeared in the top 10 lists of all three resources (**Figures 1A-C**).

**Figure 1:**
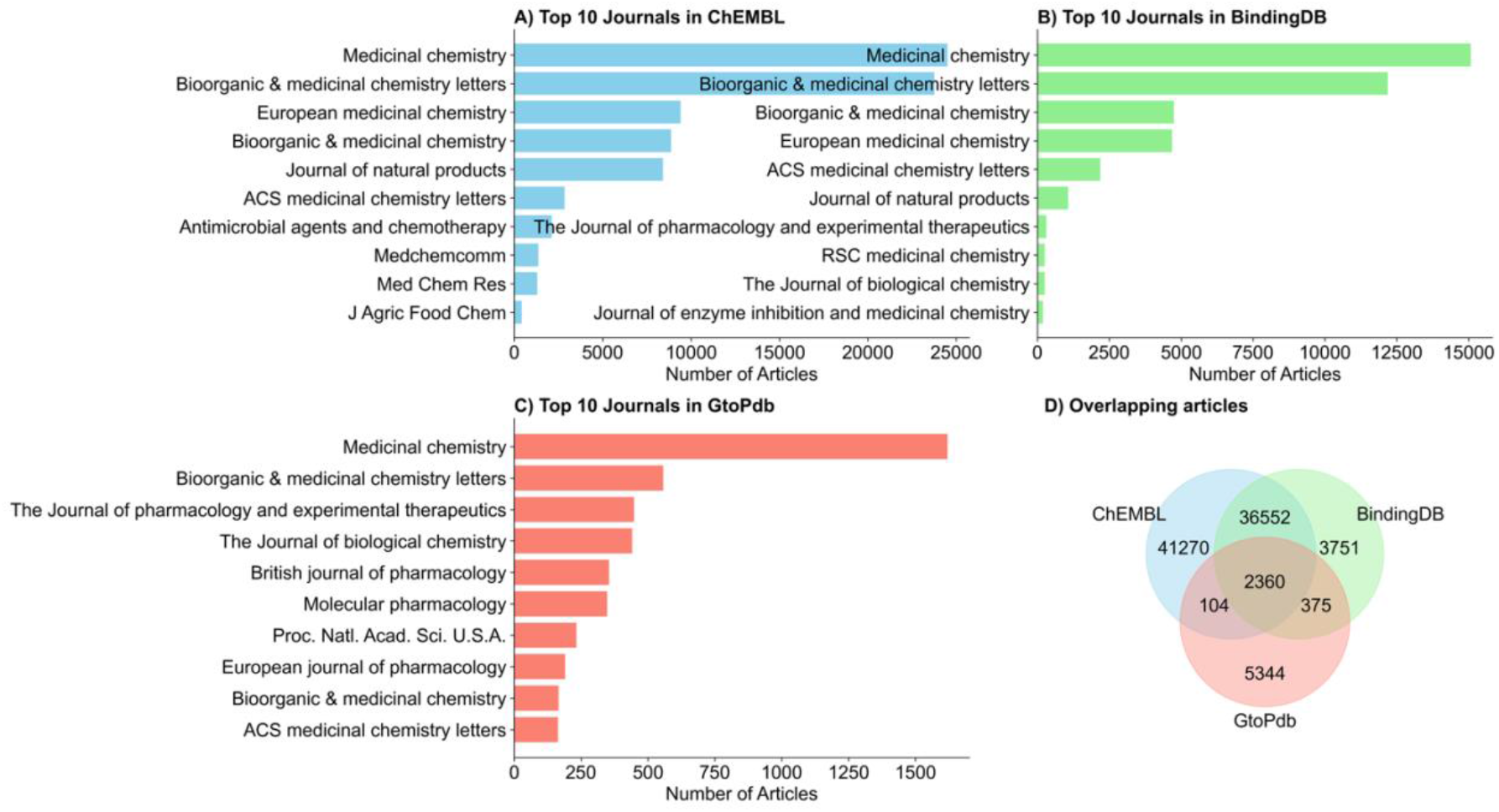
Top 10 journals contributing bioactivity data curation across three databases: **A)** ChEMBL, **B)**BindingDB, and **C)** GtoPdb. **D)** Total number of overlapping articles curated by all three databases.**-5**.

### 2.2. Preprocessing drug-target data in ChEMBL, BindingDB and GtoPdb

We downloaded the complete database dump for ChEMBL (V 33) and standalone files for BindingDB and GtoPdb to perform a comparative analysis. To enable cross-database comparisons, we standardized ligand representation using standard InChIKeys and target identifiers using UniProt IDs. In ChEMBL, ligands are annotated according to their maximum clinical development stage (e.g., approved, phase 3, phase 2, phase 1, and preclinical). For this study, we focused on approved drugs and investigational compounds (phase 1–3), excluding preclinical entries to maintain consistency with the scope of clinically relevant molecules. In the ChEMBL database dump, the max_phase column in the molecule_dictionary table indicates the highest clinical trial phase achieved by each drug-like molecule (see ChEMBL_33 schema). ChEMBL V 33 also contains a total of 7,110 compounds having max_phase > 0, comprising 3,492 approved drugs and 3,618 investigational compounds (small molecules: 6828, biological ligands: 260 and unknown:22, **Supplementary File 1, Supplementary Figure 2**). Among these, pChEMBL values, standardized negative logarithmic activity measures (‑log_10_ of molar IC_50_, EC_50_, K_ⅈ_, or Kd) are available only for 2,080 approved and 1,780 investigational compounds. To ensure consistency across analyses, we filtered out compounds lacking pChEMBL measurements. These compounds collectively bind with 2,306 single-protein targets for approved drugs and 2,187 targets for investigational compounds, representing a total of 3,067 unique protein targets. Across all bioactivity records, ChEMBL contains 104,023 binding affinity data points for compounds with max_phase > 0, corresponding to 37,651 unique drug–target pairs involving approved and investigational molecules (**Supplementary File 2**). Because multiple pChEMBL entries for the same drug–target pair often originate from independent studies employing different assay protocols or experimental conditions, we calculated the median pChEMBL value for each unique drug–target pair to ensure data consistency and comparability. All subsequent drug– target analyses for ChEMBL are performed on this refined dataset

The BindingDB dataset comprises approximately 1.4 million compounds, of which 953,851 include standard InChIKey identifiers. After excluding compounds lacking target annotations (i.e., missing UniProt identifiers), 930,213 compounds remained. Since BindingDB does not provide clinical phase information, we cross-referenced standard InChIKeys with ChEMBL to assign phase annotations. After filtering for compounds with max_phase > 0, we identified 868 approved and 642 investigational compounds, corresponding to 18,063 binding affinity measurements. Some drug–target pairs in BindingDB do not report quantitative activity values and are therefore excluded, resulting in a final subset of 774 approved and 617 investigational compounds (**Supplementary File 2**). BindingDB reports activity measurements as IC_50_, EC_50_, Kd, or K_ⅈ_ values (in nanomolar units). These are converted to pChEMBL values using the standard transformation:

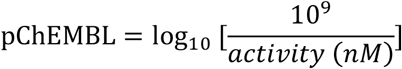

As in ChEMBL, multiple measurements for the same drug–target pair are aggregated using the median pChEMBL value. The processed BindingDB dataset includes 8,602 unique drug–target pairs, 1,510 unique compounds, and 1,410 unique protein targets (**Supplementary File 2**).

The GtoPdb database classifies compounds as either approved or non-approved. In this study, non-approved entries are assumed to represent investigational compounds. Ligand–target relationships are extracted from the GtoPdb_Compound_mapping and GtoPdb_all_interactions tables, with ligand IDs mapped to standard InChIKeys and target identifiers converted to UniProt IDs. Out of 9,979 unique ligand IDs, 6,504 are successfully mapped to standard InChIKeys, comprising 778 approved and 5,726 investigational compounds. Ligand–target pairs lacking quantitative binding data (no binding affinity values) are excluded. The final GtoPdb dataset contained 704 approved drugs, 5,726 investigational ligands, and 12,751 ligand–target pairs. As with the other datasets, when multiple pChEMBL values are reported for a given ligand–target pair, the median value is used in all subsequent analyses. **Supplementary File 2** shows all processed datasets, including both approved and investigational ligands, for ChEMBL, BindingDB and GtoPdb.

### 2.3. Classification of targets into 12 family groups for ChEMBL data

As BindingDB and GtoPdb do not offer target classification (in stand-alone files), our family-level analysis is conducted exclusively using ChEMBL’s drug-target data. ChEMBL classifies protein targets into 897 subfamilies present in the ‘Protein_classification’ table in the ChEMBL database (see ChEMBL schema).

To facilitate consistent and biologically meaningful comparisons, we manually classified ChEMBL targets into 12 high-level functional target families: enzymes (excluding kinases), kinases, G-protein-coupled receptors (GPCRs), nuclear receptors, ion channels, epigenetic regulators, adhesion receptors, transporters, cytochromes, phosphodiesterases, proteases, and an ‘other’ category for unclassified or relatively smaller families. Smaller subfamilies are grouped together under broader categories; for example, tyrosine kinases and MAP kinases are classified under ‘kinase’, while 7TM and rhodopsin-like receptors are grouped under ‘GPCR’. Conversely, in some cases, subgroups are separated to enhance resolution and interpretability, for instance, kinases are treated as distinct from the broader enzyme class, and adhesion receptors, typically considered a GPCR subclass, are assigned their own category.

Figure 2 illustrates the distribution of ChEMBL targets, drugs, and interactions for approved and investigational compounds across these 12 target families, without applying any potency cutoff on pChEMBL values. A more stringent version of this analysis, using a pChEMBL of ≥ 5, pChEMBL of ≥ 6 and pChEMBL of ≥ 7 is presented respectively in **Supplementary Figures 3-5**. Among the 12 target families, enzymes, kinases, and GPCRs exhibited the broadest binding coverage for both approved and investigational compounds. The complete mapping of 3083 ChEMBL targets to their assigned high-level families is provided in **Supplementary File 3**.

**Figure 2:**
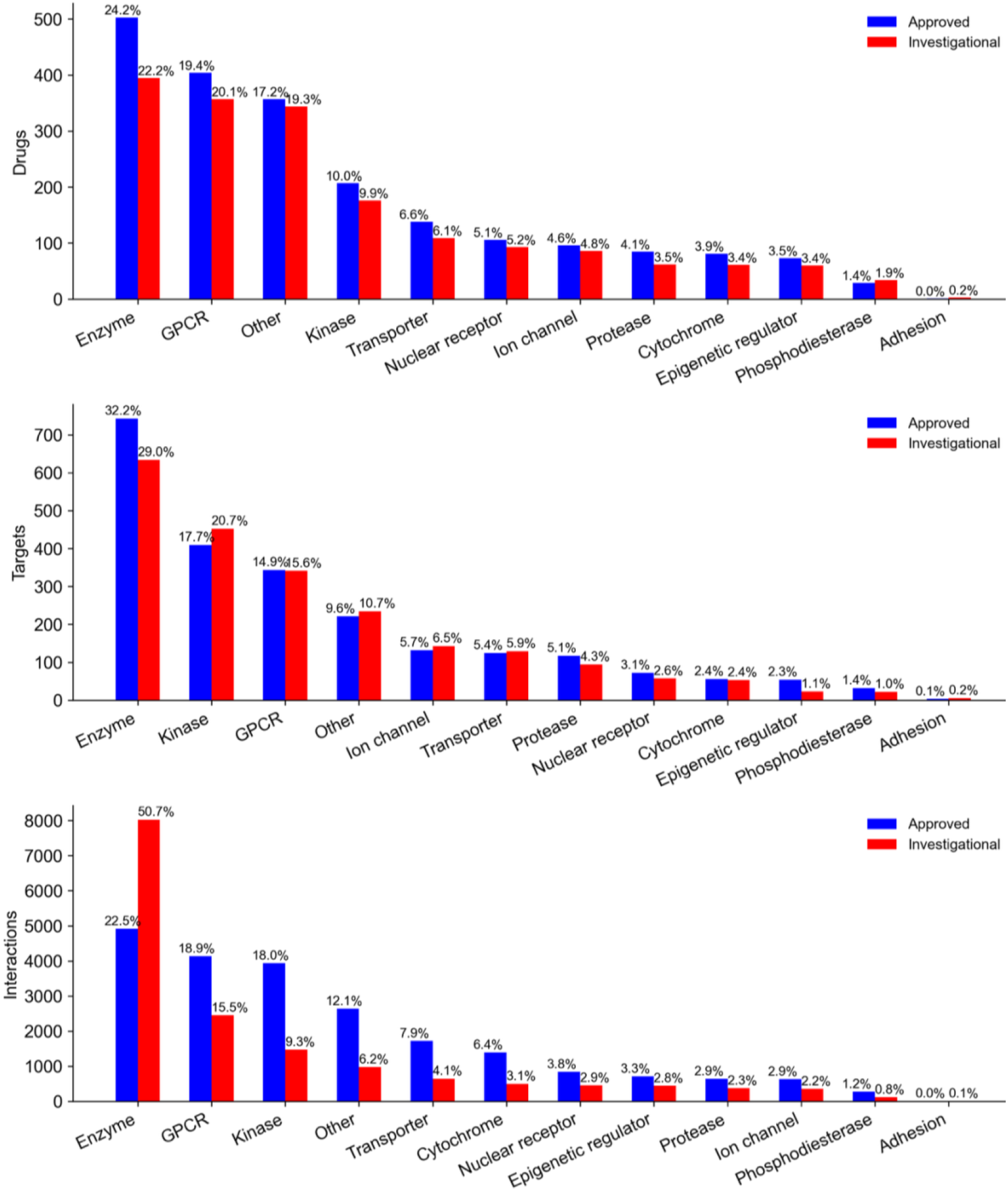
Frequencies of drugs, targets, and their interactions are shown across 12 target families associated with approved drugs (blue) and investigational compounds (red) in the ChEMBL database. Target families with limited representation or undefined classification are grouped under the ‘Other’ category. The percentage values above each bar indicate the proportion of each target class within the set of approved drugs and investigational compounds. An activity threshold of pChEMBL ≥ 0 was applied for inclusion in this analysis. A more stringent version of this analysis, using a pChEMBL of ≥ 5, pChEMBL of ≥ 6 and pChEMBL of ≥ 7 is presented respectively in **Supplementary Figures 3-5**.

### 2.4. Classification of indications into families

ChEMBL (V 33) links 2,330 approved drugs to 817 clinical indications, manually curated from reputable sources including ClinicalTrials.gov, DailyMed, the FDA, EMA, and others. This drug–indication mapping is accessible via the ‘Drug_indication’ table in ChEMBL schema. To support higher-level pharmacological interpretation and cross-disease analyses, we manually grouped these 817 indications into 28 broad disease categories. These include infections, cardiovascular, mental, neoplasms (non cancerous), respiratory, cancer, nervous system, musculoskeletal, digestive system, eye, nutritional and metabolic, skin and connective tissue, hematologic, immunological, urogenital diseases, pathological conditions, dermatologic, renal, reproductive system, parasitic, ear, genetic and congenital, dental, endocrine system, head and neck, oral diseases, biliary tract and sleep disorders indication groups. **Supplementary File 9** presents the comprehensive mapping of 817 indications into 28 indication groups, along with their associated 2,330 approved drugs and supporting reference links.

Figure 3 illustrates the distribution of approved drugs across the indication groups, with six smaller categories combined under “Other.” The infection indication group accounts for the largest number of approved drugs (n=751), followed by cardiovascular-related indications (n=666). Since some drugs are approved for multiple indications, overlaps exist across indication groups, which is further visualized via a chord diagram in **Figure 6**. Together, this structured classification offers a practical resource for future research in drug repurposing, indication-based compound prioritization, and pharmacological landscape modeling.

**Figure 3:**
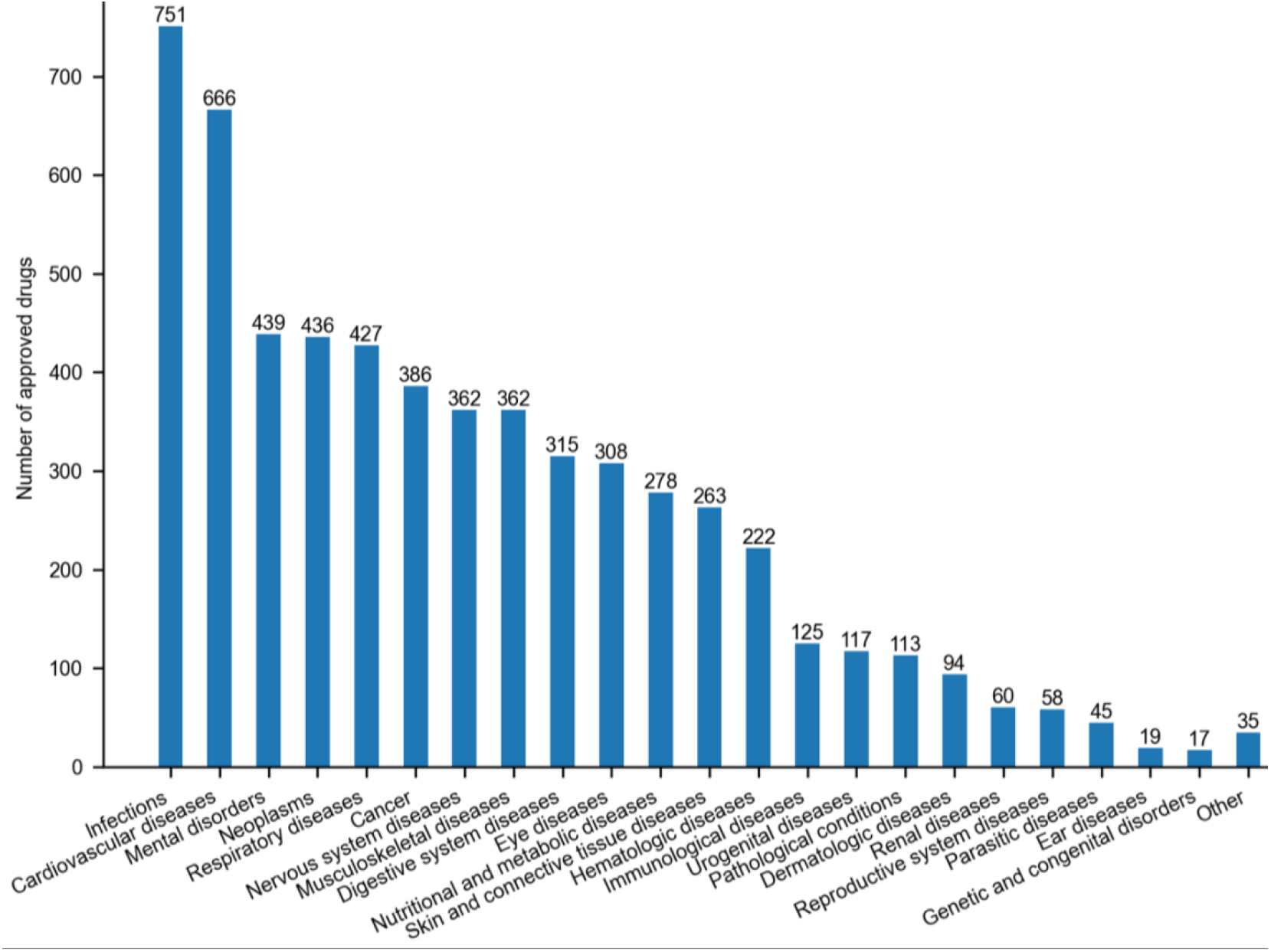
Number of drugs approved across different indication groups (smaller groups are combined under ‘Other’ category)

### 2.5. Comparative analysis of drug target data in ChEMBL, BindingDB and GtoPdb

To enable a comparative analysis across the three drug–target databases, ChEMBL, BindingDB, and GtoPdb, we first examined target coverage for both approved and investigational compounds. A series of word cloud visualizations illustrate the top 50 approved drugs represented across the three databases at four activity thresholds: pChEMBL ≥ 0 (**Supplementary Figure 6**), pChEMBL ≥ 5 (**Supplementary Figure 7**), pChEMBL ≥ 6 (**Supplementary Figure 8**), and pChEMBL ≥7 (**Supplementary Figure 9**). The font size of each drug name reflects the number of protein targets associated with that drug (bigger font size represents more targets). At pChEMBL ≥ 0, sunitinib is the most broadly profiled drug in both ChEMBL (290 targets) and BindingDB (178 targets), while flufenamic acid ranked highest in GtoPdb with 33 targets.

Without applying any threshold on binding affinity (i.e., pChEMBL ≥ 0), the average number of targets per approved drug is highest in ChEMBL (mean = 10), followed by BindingDB (mean = 7), and GtoPdb (mean = 3), as shown in **Figure 4**. A similar trend is observed for investigational compounds, with mean target counts of 9 in ChEMBL, 5 in BindingDB, and 2 in GtoPdb. These findings suggest that ChEMBL provides the broadest proteome-level target coverage among the three databases. This pattern remained consistent under more stringent activity thresholds: pChEMBL ≥5 (**Supplementary Figure 10**), pChEMBL ≥6 (**Supplementary Figure 11**), and pChEMBL ≥7 (**Supplementary Figure 12**), demonstrating the robustness of ChEMBL’s target profiling across confidence levels.

**Figure 4:**
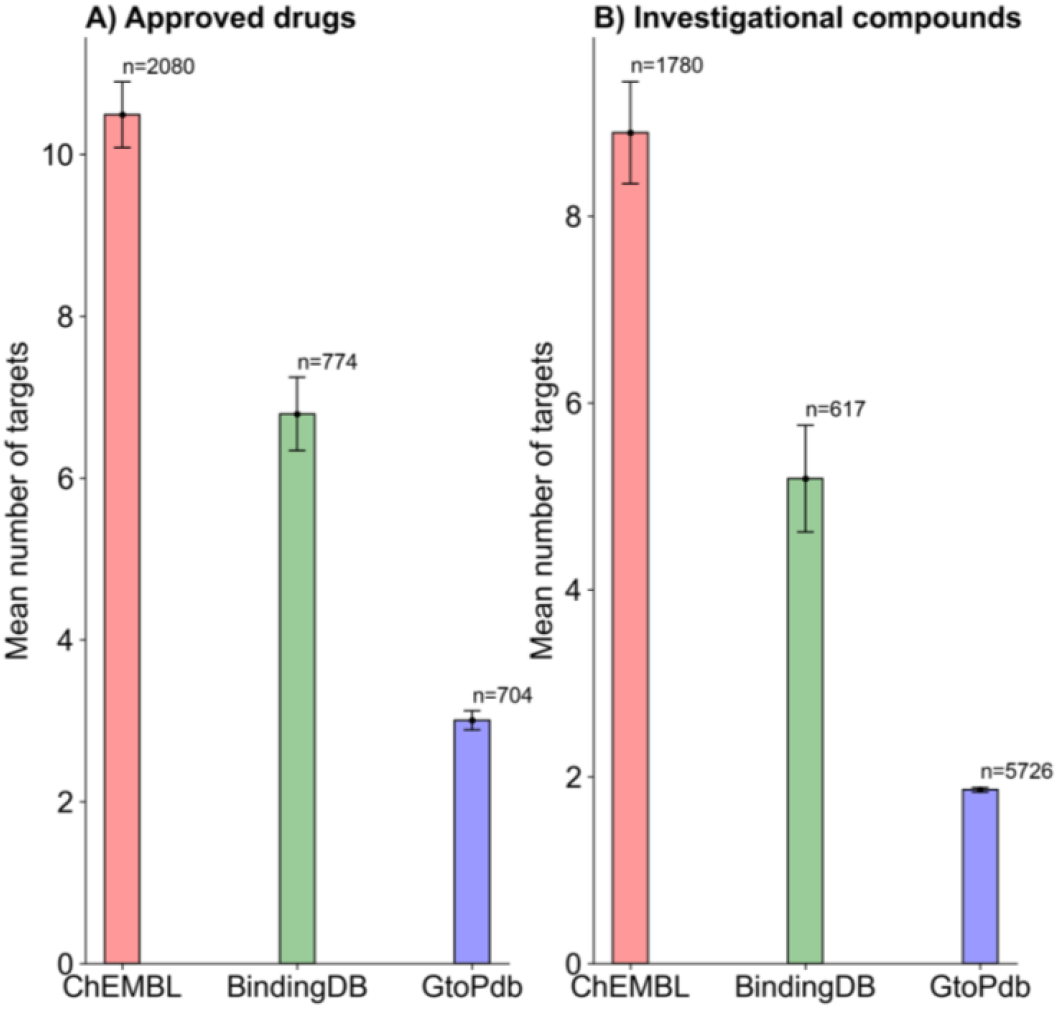
Mean number of targets across three databases at a pChEMBL ≥ 0. **A)** Approved drugs, **B)** Investigational compounds. ‘n’ indicates the total number of drugs or compounds included in each analysis. A more stringent version of this analysis, using a pChEMBL of ≥ 5, pChEMBL of ≥ 6 and pChEMBL of ≥ 7 is presented respectively in Supplementary Figures 10-12.

We further evaluated the overlap in drug, target, and interaction data across the three databases for both approved and investigational compounds. Among approved drug– target pairs, 371 drugs, 283 targets, and 535 interactions are found to be common across all three databases (**Figure 5**; pChEMBL ≥ 0). For investigational compounds, 255 compounds, 521 targets, and 316 interactions overlapped. Drug–target overlaps for both approved and investigational compounds at three additional activity thresholds: pChEMBL ≥ 5, pChEMBL ≥ 6, and pChEMBL ≥ 7, are presented in **Supplementary Figures 13–15**. Overall, ChEMBL exhibited the most comprehensive proteome-scale annotation, consistently surpassing BindingDB and GtoPdb in both the breadth and depth of its drug–target interaction coverage.

**Figure 5:**
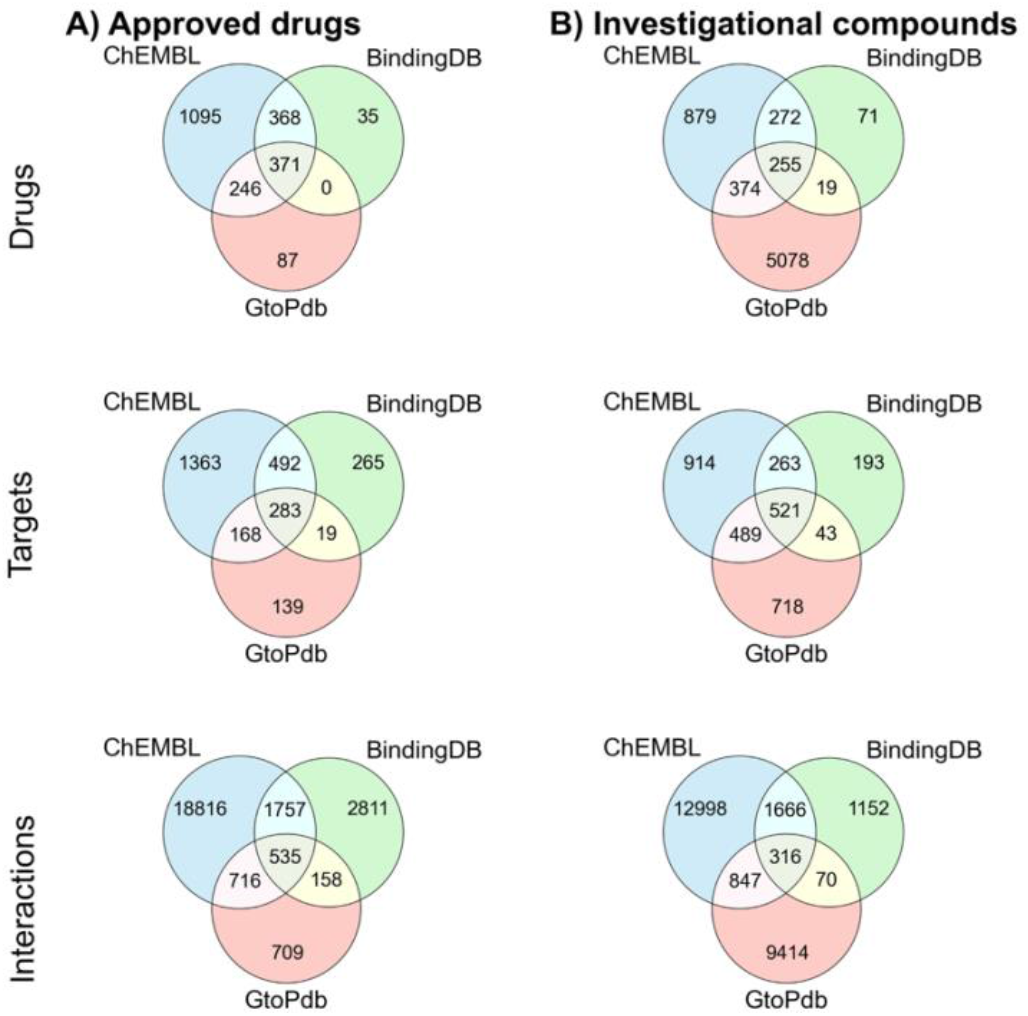
Overlap of drugs, targets, and interactions across approved drugs and investigational compounds in the three binding affinity databases. Target overlap is determined using UniProt IDs, and compound overlap is based on standard InChIKey identifiers. Drug–target overlaps for both approved and investigational compounds at three additional activity thresholds: pChEMBL ≥5, pChEMBL ≥6, and pChEMBL ≥7, are presented in **Supplementary Figures 13–15**.

### 2.6. Physiochemical properties

To better understand the chemistry associated with approved drugs, we extracted 15 physicochemical properties from the ChEMBL database. These properties include: alogP, cx_logD, cx_logP, cx_most_apKa, cx_most_bpKa, full_mwt, hba, hba_lipinski, hbd, hbd_lipinski, mw_freebase, np_likeness_score, psa, qed_weighted, and rtb. Detailed definitions and computational methodologies for these descriptors are available in ChEMBL’s Compound_Properties table (ChEMBL Schema). **Supplementary File 4** provides physicochemical properties information for all approved drugs across 28 indications groups. Supplementary information further provides a section on analysis of drug’s likeliness properties for approved drugs and investigational drugs present in ChEMBL database (**Supplementary Figure 24-26**).

## 3. RESULTS AND DISCUSSION

### 3.1. Indications groups with shared drug repurposing potential

Section 2.4 describes the categorization of 817 therapeutic indications from the ChEMBL database into 28 broad disease groups, highlighting the number of approved drugs associated with each group (**Figure 3**). **Supplementary File 9** provides detailed information on these drugs, including the original ChEMBL indication names (column: Indiction_mesh_name), the corresponding high-level disease group classifications annotated in this study (column: Indication_group), and the supporting web links documenting these associations. In this section, we investigate the extent of drug sharing across these indication groups, an analysis that may offer insights into therapeutic overlaps and repurposing potential. **Figure 6** provides a visual summary of drug overlap between indication groups, where the width of each connecting chord represents the number of shared approved drugs. While this figure conveys the relative scale of overlap, the exact numbers are detailed in **Supplementary File 5**. For example, the cancer group is associated with 386 approved drugs, 103 of which are also approved for neoplasms, 30 for hematologic diseases, and 15 for nutritional and metabolic disorders. These overlaps reflect well-established biological relationships. For example, hematologic malignancies and solid tumors may share key molecular targets, most notably tyrosine kinases such as BCR‑ABL and KIT, which can be targeted by the same drugs across multiple cancer types. A prominent example is imatinib, originally developed for chronic myeloid leukemia (CML) via BCR‑ABL inhibition, and later approved for gastrointestinal stromal tumors (GISTs) through its inhibition of KIT and PDGFR [23, 24]. Furthermore, metabolic reprogramming is a recognized hallmark of cancer and is observed across both hematologic and solid malignancies [25, 26] This commonality supports the observed therapeutic overlap across cancer, hematologic, and metabolic disease indications. The largest indication group, infections (n = 751 approved drugs), also shows substantial overlap, most notably with respiratory (n = 67 shared drugs) and eye diseases (n = 55). This overlap is biologically plausible: many respiratory pathogens such as adenoviruses, haemophilus influenzae, streptococcus pneumoniae, and moraxella catarrhalis can infect both the lungs and the ocular surface, causing conjunctivitis, keratitis, and pneumonia [27]. These conditions caused by bacterial infections are often managed using shared classes of antimicrobials and anti-inflammatory agents [28].

This overlap analysis reveals disease groups sharing pharmacologically active compounds, such as broad-spectrum agents targeting conserved mechanisms across pathogens [29], providing a rational basis for drug repurposing. These insights can guide candidate prioritization and cross-indication screening, accelerating therapy development for related conditions.

**Figure 6:**
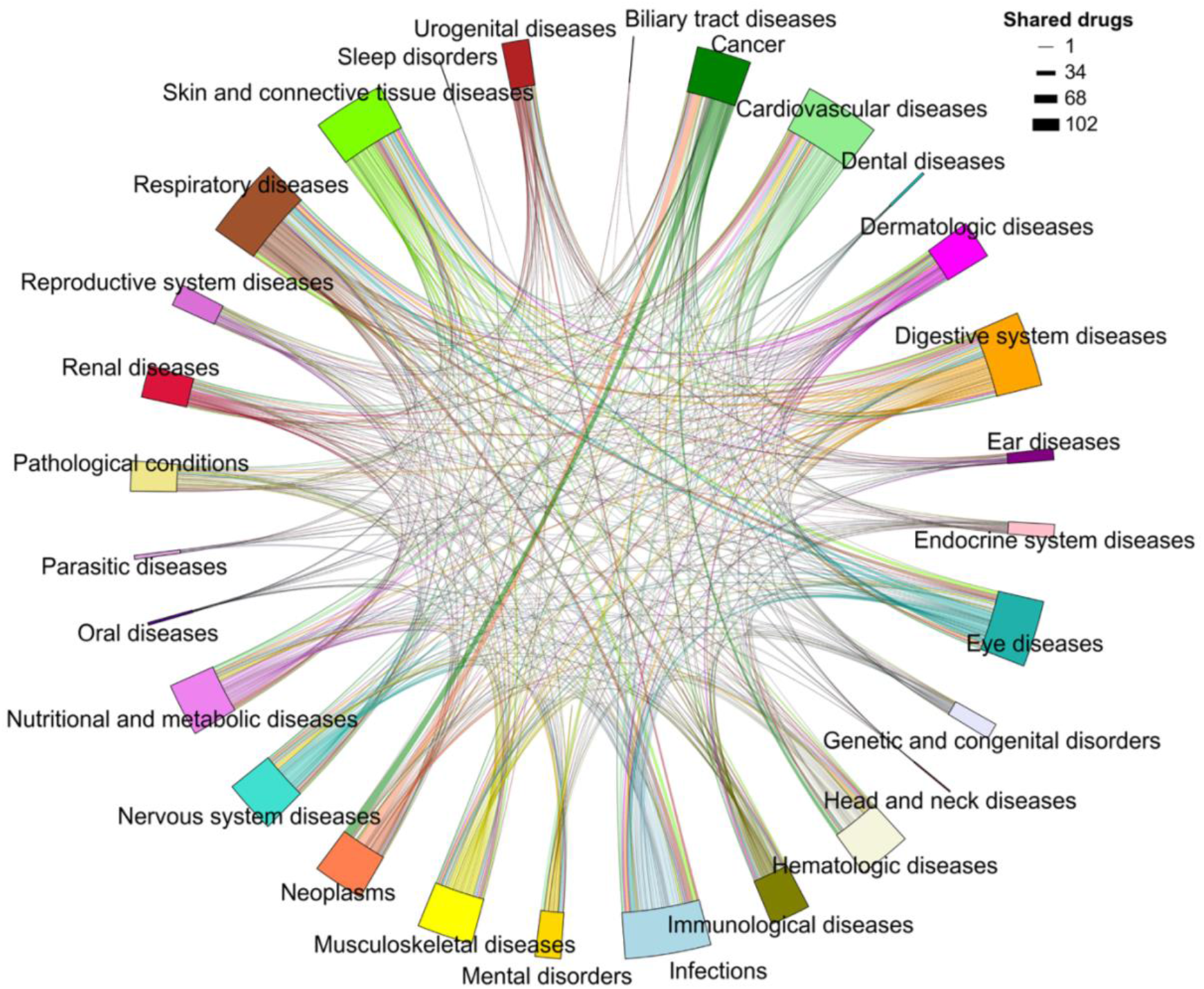
The number of drugs shared among 28 indication groups is represented, with the width of each connecting patch proportional to the number of shared approved drugs. **Supplementary File 5** presents this information in a tabular format.

### 3.2. Associations between physiochemical properties and indications groups

Previously, we categorized approved ChEMBL drugs into 28 therapeutic indication groups based on their annotated clinical uses (**Supplementary File 9**). Building on this classification, we next examined whether distinct physicochemical property patterns could be identified among approved drugs across these indication groups. **Supplementary Figures 16–19** illustrate the distribution of physicochemical properties across all 28 groups. Using these data, we defined upper and lower threshold values for each property within every indication group, calculated as ±5% around the mean value observed for drugs in that group (**Supplementary File 6**). This property-based profiling establishes indication-specific physicochemical boundaries, offering practical guidance for early-stage drug design and optimization. For instance, drugs approved for nervous system disorders typically display lower polar surface area (PSA) and fewer hydrogen bond donors (HBD), consistent with the need for central nervous system penetration [30]. Our analysis confirms this trend, showing PSA values ranging from 54.65 to 60.41, the second lowest among the 28 groups and HBD values between 0.95 and 1.05, the lowest observed (**Supplementary File 6**). Collectively, this analysis provides a systematic framework linking physicochemical profiles to therapeutic groups, supporting the rational design of compounds more likely to succeed in specific indications.

Because different physicochemical properties span varying numerical ranges, we normalized all 15 properties to z-scores to enable direct comparison and clustering of indication groups based on the physicochemical characteristics of approved drugs. For indication groups with multiple approved drugs, we took the mean of z-scores for each property, yielding a single representative value per property across the 28 indication groups. Positive z-scores indicate above-average physicochemical property values, whereas negative z-scores indicate below-average values across indication groups. Hierarchical clustering of indication groups and physicochemical properties revealed distinct and interpretable patterns; for example, drugs approved for dental diseases and head and neck disorders exhibited notably similar physicochemical profiles (**Figure 7**). Likewise, drugs approved for cancers and neoplasms displayed overlapping physicochemical characteristics, reflecting shared molecular design principles across oncology indications. Overall, this systematic analysis highlights how physicochemical property profiling can inform indication-specific molecule design and guide future drug development efforts for unmet medical needs.

**Figure 7:**
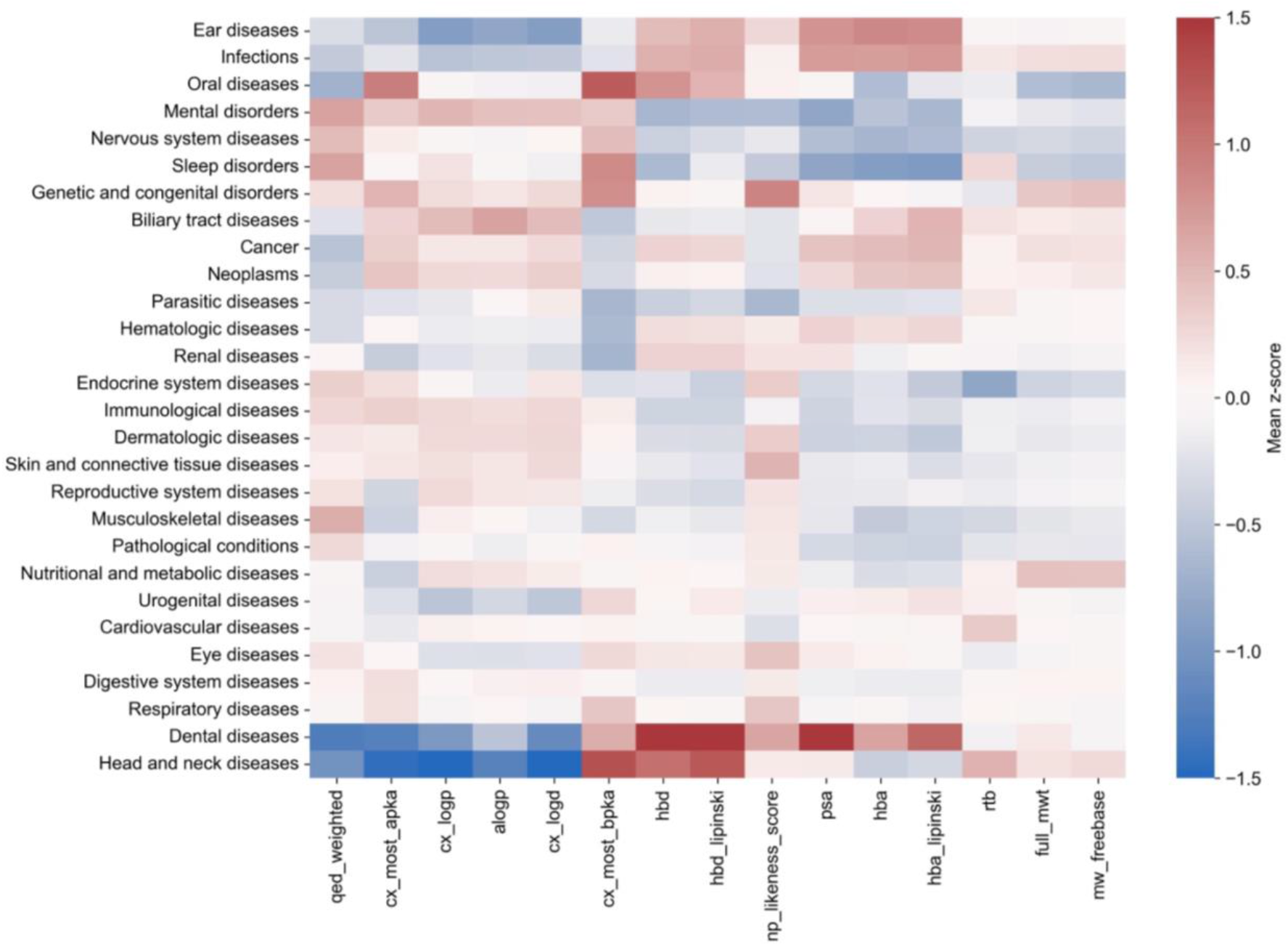
Mean z-scores of 15 physiochemical properties for the approved drugs across 28 indications groups. Z-scores are computed as each physiochemical property has different value ranges. Positive z-score values indicate higher-than-average property values; negative values indicate lower-than-average values across indication groups.

### 3.3. Top 10 drugs approved for most indications

We conducted a systematic analysis of the top repurposed drugs across diverse disease indications. The ten most frequently repurposed drugs identified are: dexamethasone, prednisone, prednisolone, cortisone acetate, menthol, naproxen, lidocaine, methotrexate, doxorubicin hydrochloride, and ibuprofen (**Table 1**). These drugs demonstrate substantial versatility in their therapeutic applications. Notably, dexamethasone is a synthetic glucocorticoid that primarily exerts its effects through the glucocorticoid receptor (UniProt ID: P04150), a ligand-activated transcription factor. It received FDA approval in 1958 and is subsequently introduced into clinical use for hematologic malignancies, including acute lymphoblastic leukemia (ALL) [31, 32]. Since its approval, dexamethasone is repurposed for 74 distinct indications spanning a broad range of indication groups, including oncology, respiratory diseases, endocrinology, dermatology, and autoimmune disorders [33] (**Table 1, Supplementary File 7**). Mechanistically, dexamethasone binds to cytosolic glucocorticoid receptors and translocates to the nucleus, modulating gene transcription involved in inflammatory, immune, metabolic, and apoptotic pathways, serving as a potent ligand-activated transcription factor [34]. ChEMBL data indicate interactions with 11 distinct protein targets, underscoring its pharmacological flexibility (**Table 1**).

**Table 1:** Top 10 repurposed drugs showing number of associated indications and targets. Supplementary File 7 shows weblinks for evidence of repurposing of each of the 10 drugs.

| Drug | Indications | Targets | Target families |
| --- | --- | --- | --- |
| Dexamethasone | 74 | 11 | Nuclear receptor, G protein-coupled receptors, Cytochrome, Other |
| Prednisone | 62 | 7 | Enzyme, Nuclear receptor, Other, Phosphodiesterase |
| Prednisolone | 58 | 13 | Enzyme, G protein-coupled receptors, Nuclear receptor, Other, Protease, Transporter |
| Cortisone acetate | 44 | 3 | Enzyme, G protein-coupled receptors, Nuclear receptor |
| Menthol | 30 | 4 | Ion channel |
| Lidocaine | 25 | 11 | Cytochrome, Enzyme, G protein-coupled receptors, Ion channel, Kinase, Other |
| Naproxen | 24 | 14 | Cytochrome, Enzyme, G protein-coupled receptors, Transporter |
| Methotrexate | 22 | 50 | Enzyme, Epigenetic regulator, Other, Phosphodiesterase, Protease, Transporter |
| Ibuprofen | 22 | 24 | Cytochrome, Enzyme, Other, Transporter |
| Doxorubicin hydrochloride | 22 | 35 | Cytochrome, Enzyme, Epigenetic regulator, G protein-coupled receptors, Kinase, Nuclear receptor, Other, Phosphodiesterase, Protease, Transporter |

Interestingly, 7 of the 10 most frequently repurposed drugs in our analysis interact with more than 10 distinct protein targets, highlighting polypharmacology as a common trait among highly repurposable therapeutics. This aligns with broader evidence that multi-target activity is a hallmark of efficacious agents used across multiple indications [35]. **Table 1** summarizes the top 10 drugs along with associated indications, targets, and target superfamilies. A more detailed version of this table is presented in **Supplementary File 7**.

### 3.5. A novel pathway driven strategy to repurpose drugs for 10 cancer types

To further guide, how publically available datasets can help to design drug repurposing pipelines, we implemented here pathway-based reference pipeline to identify repurposed drugs for ten cancer types: prostate, endometrial, breast, non-small cell lung (NSCLC), colorectal, pancreatic, hepatocellular cancer (HCC), gastric, thyroid, and acute myeloid leukemia (AML). These cancer types are chosen due to the availability of well-curated and disease-specific pathway information in the Kyoto Encyclopedia of Genes and Genomes (KEGG) database [36], enabling a biologically grounded approach to target identification. For each cancer type, we retrieved the KEGG disease ID, MeSH ID, and the associated KEGG disease pathways. From these pathways, we extracted gene symbols functionally implicated in the disease mechanisms. These gene symbols are then mapped to UniProt identifiers using the UniProt API (reviewed entries), providing a standardized and biologically relevant protein target set. A total of 377 unique genes are associated with the ten cancer types, of which 375 are successfully mapped to UniProt IDs. The number of pathway-associated protein targets identified for each cancer type is thyroid (37), NSCLC (72), AML (67), endometrial (58), pancreatic (76), prostate (105), colorectal (86), breast (147), gastric (148), and HCC (168). **Supplementary Table 1** shows names of gene targets involved in disease pathways in the KEGG database.

To identify repurposing candidates, we first retrieved drug–target bioactivity data from ChEMBL (V 33), filtering for approved drugs with known activity against single human protein targets and available pChEMBL values. For each cancer type, potent approved drugs targeting a good number of overlapping number of pathway associated targets can be repurposing candidates for that cancer type. As there is no such consensus on potency cut-offs, drug-target bioactivity analysis for this pipeline is performed on 3 different potency cut-offs (pChEMBL ≥5, pChEMBL ≥6 and pChEMBL ≥7). For each threshold, when multiple bioactivity measurements exist for a given drug–target pair across different studies, the median pChEMBL value is computed to ensure data robustness. **Figure 8** presents the sensitivity analysis used to identify repurposing candidates across ten cancer types at the three potency cut-offs. As expected, a greater number of candidates are identified at lower potency thresholds, while stricter cut-offs resulted in fewer, but potentially more specific, candidates. Previous studies have suggested that a ≥10% overlap between drug-binding targets and pathway-associated targets can indicate meaningful pathway-level association [37]. Based on this criterion, we further analyzed repurposing candidates exhibiting ≥10% pathway target overlap for ten cancer types at pChEMBL ≥5 (**Supplementary Figure 20**), pChEMBL ≥6 (**Supplementary Figure 21**), and pChEMBL ≥7 (**Supplementary Figure 22**). **Supplementary Tables 2 and 3** list the repurposed drugs and their corresponding target proteins involved in cancer-specific pathways for pathway coverage ≥10% at potency thresholds of pChEMBL ≥ 6 and pChEMBL ≥ 5, respectively.

**Figure 8:**
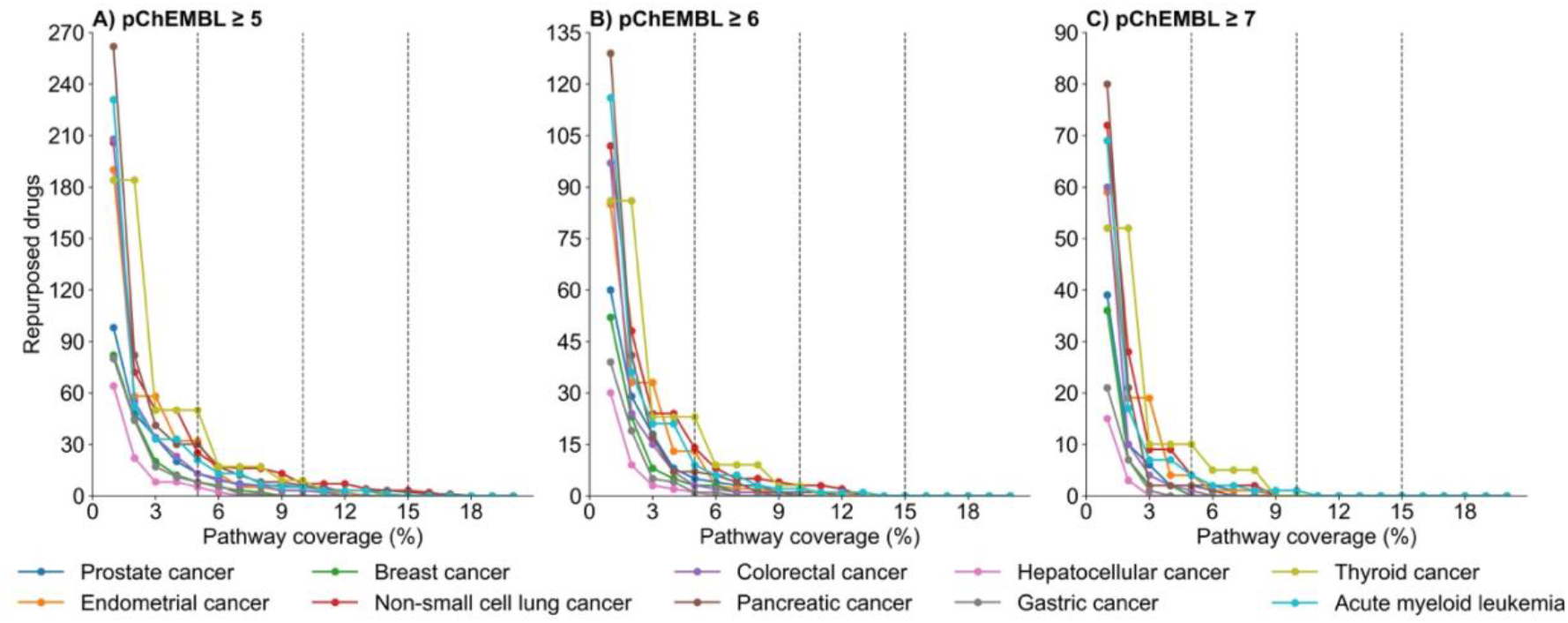
Number of repurposing candidates (Y-axis) identified across ten cancer types at three potency thresholds: **(A)** pChEMBL ≥ 5, **(B)** pChEMBL ≥ 6, and **(C)** pChEMBL ≥ 7. The X-axis represents the percentage of pathway targets overlapped by the binding targets of repurposing candidate drugs.

These results illustrate the potential of leveraging pathway-level biological context to systematically uncover repurposing therapeutic opportunities. For analysis in **Figure 8 and Supplementary Figures 20-22**, we excluded from consideration any drug already approved for the same cancer type. This is done by cross-referencing each candidate against known approved indications as listed in **Supplementary File 4. Supplementary Figure 23** presents the known number of approved drugs for each cancer type, while **Supplementary Table 4** lists their corresponding names. Of the ten cancers analyzed, breast cancer is associated with the largest number of approved drugs (n=31, **Supplementary Figure 23**), while endometrial, gastric, and hepatocellular carcinoma (HCC) have no approved therapies recorded in the ChEMBL database. This case study introduces a scalable, pathway-informed drug repurposing pipeline that integrates curated biological knowledge and quantitative drug-target interaction data. Although demonstrated here across ten cancer types, the reference pipeline is applicable to any other indication for which pathway information is available. By targeting biologically validated networks rather than isolated proteins, this strategy increases the likelihood of identifying repurposing candidates with mechanistic relevance and therapeutic efficacy [38].

## 4. CONCLUSION

This study presents a systematic, data-driven methodologies for drug repurposing that integrates high-quality drug–target data, physicochemical properties-based clustering of indication groups, and a reference pathway-driven pipeline. By leveraging the most comprehensive DTI databases, ChEMBL, BindingDB, and GtoPdb, we provide an in-depth comparative analysis of their curation practices, coverage, and binding data (see **Figures 1, 2, 4**, and **5**). Hierarchical clustering of indication groups and physicochemical properties revealed distinct and interpretable patterns among drugs approved for different indications (**Figure 7, Supplementary Figures 16–19 and Supplementary File 6**). These findings can guide compound prioritization during early-stage drug discovery by aligning candidate properties with those historically successful within each therapeutic area. This approach enhances the efficiency of medicinal chemistry efforts, though it may also oversimplify pharmacological complexity and overlook emerging novel chemotypes or drug modalities that deviate from established norms [39].

The pathway-driven use case (Section 3.5), applied to ten major cancer types, demonstrates how curated biological networks can be systematically leveraged to identify promising repurposing candidates based on mechanistic relevance and target coverage (**Figure 8; Supplementary Figures 20–23**). The strength of this work lies in its structuring of drug–target data, physicochemical profiles, and indication-specific context into a unified framework. By connecting these curated datasets to pathway-level and mechanistic insights, our approach enables more precise identification and prioritization of repurposing opportunities with strong biological plausibility.

Several limitations should be acknowledged. Although some rare diseases are included among the 817 indications (Supplementary File 9), they were not categorized as a separate group. For instance, Tourette syndrome, a rare neurological disorder, was grouped under nervous system diseases, while Gaucher disease and Wilson disease were assigned to the nutritional and metabolic diseases category. Another limitation relates to the reliance on KEGG pathways, which provide structured representations of disease biology but may not fully capture tissue- or patient-specific molecular variation.

Additionally, our analysis focused primarily on pharmacodynamic relationships, without incorporating pharmacokinetic parameters, target expression, or toxicity data, all of which are critical for translational applications [38]. Data availability further constrained the scope to small-molecule drugs, with limited representation from other therapeutic modalities such as peptides and proteins. Future extensions could integrate transcriptomic, proteomic, and clinical outcome datasets to enable context-aware precision medicine [40], as well as incorporate adverse event profiles, drug-induced gene expression signatures [41], and network-based proximity metrics to refine prioritization, as demonstrated in prior studies [42]. Applying the pathway-driven repurposing framework beyond oncology to rare, neglected, or complex diseases, could further expand its clinical and translational utility [43].

## CONFLICTS OF INTEREST

The authors declare no conflicts of interest.

## ACKNOWLEDGMENTS

The work was funded by the Research Council of Finland (No. 351507 to ZT).

Susanna Savander is a Master’s student at the Institute for Molecular Medicine Finland (FIMM), HiLIFE, University of Helsinki, Finland.

Nurettin Nusret Curabaz is an Erasmus student at FIMM, HiLIFE, University of Helsinki, Finland.

Amna Mumtaz Abbasi is visiting research assistant at FIMM, HiLIFE, University of Helsinki, Finland.

Asifullah Khan is professor at Pakistan Institute of Engineering & Applied Sciences (PIEAS), Islamabad, Pakistan

Khalid Saeed is an associate scientist at King Abdullah International Medical Research Center (KAIMRC), King Saud bin Abdulaziz University for Health Sciences.

Ziaurrehman Tanoli is a Principal Investigator at FIMM, HiLIFE, University of Helsinki, Finland.

## Notes

### Competing Interest Statement

The authors have declared no competing interest.

### Summary of Updates

. In summary we updated several figures and tables, added 12 new supplementary and 2 new supplementary files. These revisions further enhanced the clarity and overall quality of the manuscript.

