## Supplementary information for "Data-driven strategies for drug repurposing: insights, recommendations, and case studies"

**SUPPLEMENTARY MATERIAL**

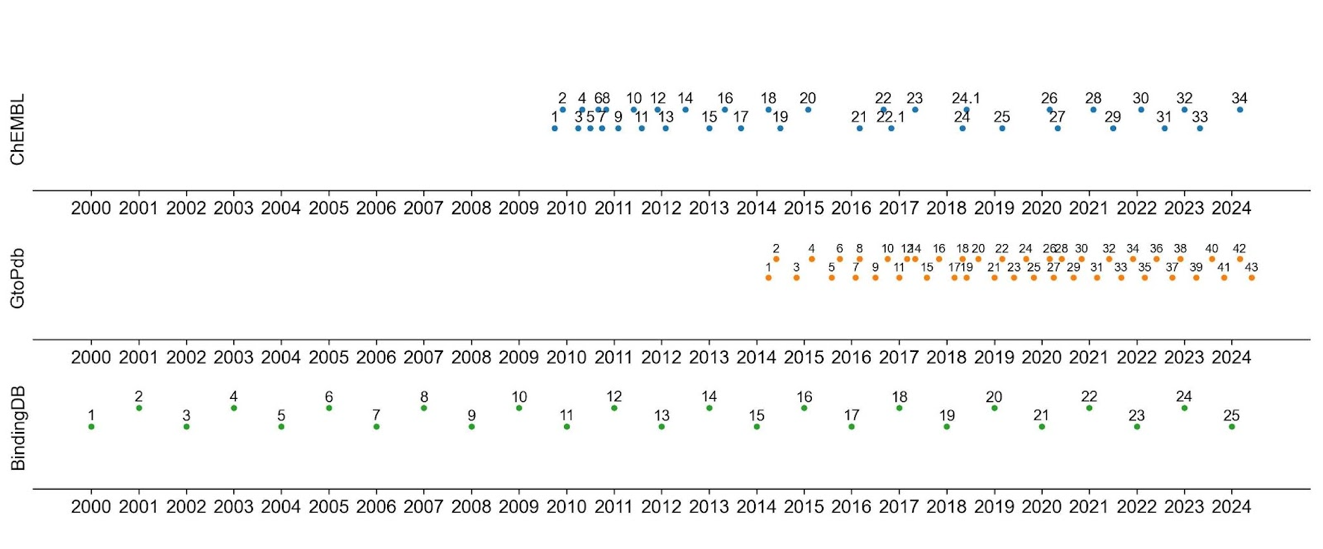

**Supplementary Figure 1:** The evolution and timeline for public releases of ChEMBL, GtoPdb, and BindingDB from their inception to the present.

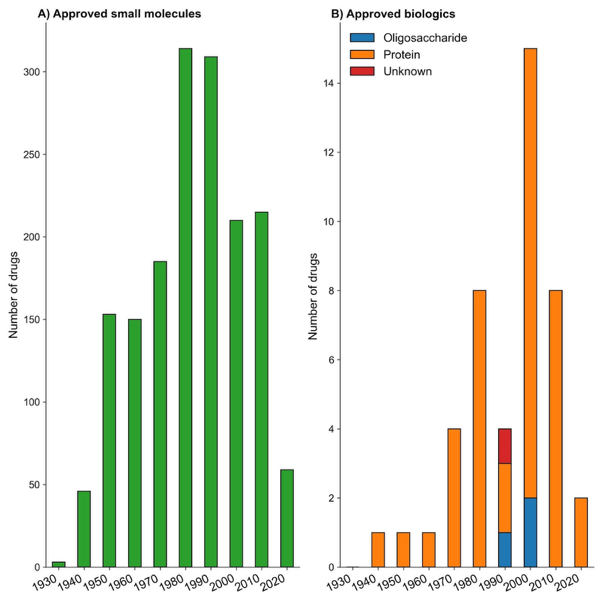

**Supplementary Figure 2:** Approved drugs by decade for the past century based on the data from the ChEMBL database. The x-axis describes the year of first approval of each drug. Each drug is represented only once, regardless of any subsequent approvals for additional indications. Small molecules and other biologics are displayed in separate plots as the number of approved small molecules is significantly higher.

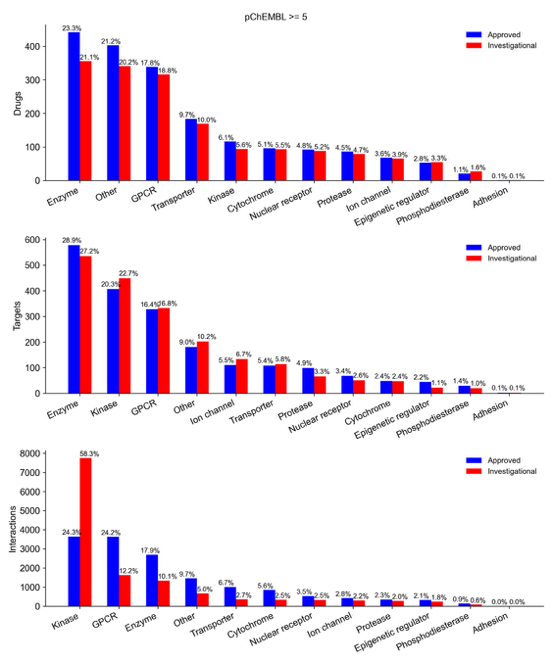

**Supplementary Figure 3:** Frequencies of drugs, targets, and their interactions are shown across 12 target families associated with approved drugs (blue) and investigational compounds (red) in the ChEMBL database. Target families with limited representation or undefined classification are grouped under the 'Other' category. The percentage values above each bar indicate the relative proportion of each target class within the set of approved drugs or investigational compounds. An activity threshold of pChEMBL ≥5 was applied for inclusion in this analysis.

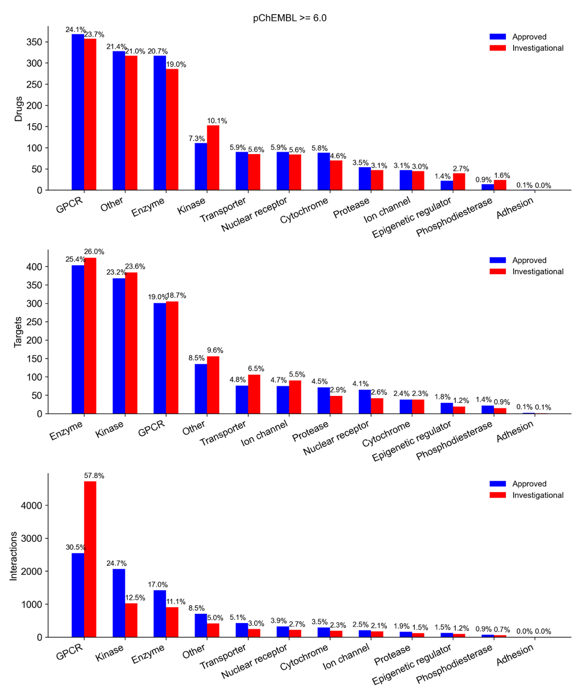

**Supplementary Figure 4:** Frequencies of drugs, targets, and their interactions are shown across 12 target families associated with approved drugs (blue) and investigational compounds (red) in the ChEMBL database. Target families with limited representation or undefined classification are grouped under the 'Other' category. The percentage values above each bar indicate the relative proportion of each target class within the set of approved drugs or investigational compounds. An activity threshold of pChEMBL ≥6 was applied for inclusion in this analysis.

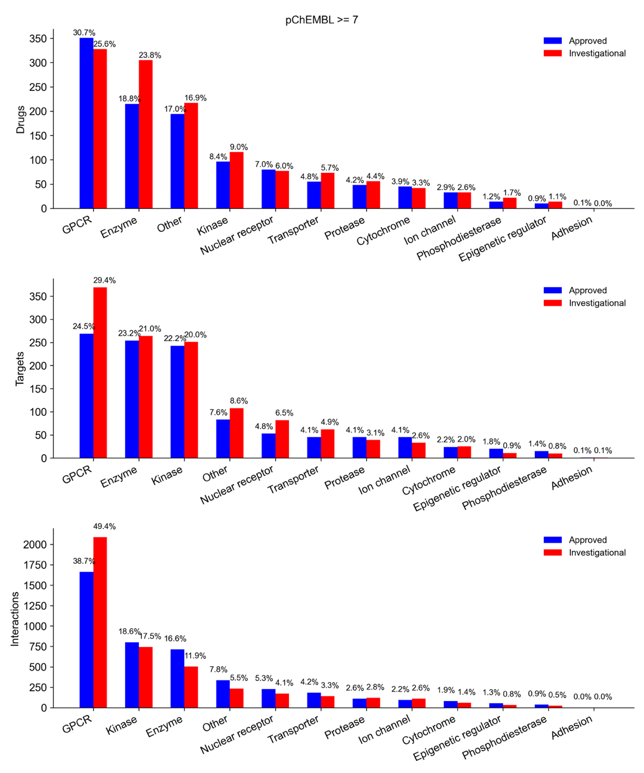

**Supplementary Figure 5:** Frequencies of drugs, targets, and their interactions are shown across 12 target families associated with approved drugs (blue) and investigational compounds (red) in the ChEMBL database. Target families with limited representation or undefined classification are grouped under the 'Other' category. The percentage values above each bar indicate the relative proportion of each target class within the set of approved drugs or investigational compounds. An activity threshold of pChEMBL ≥7 was applied for inclusion in this analysis.

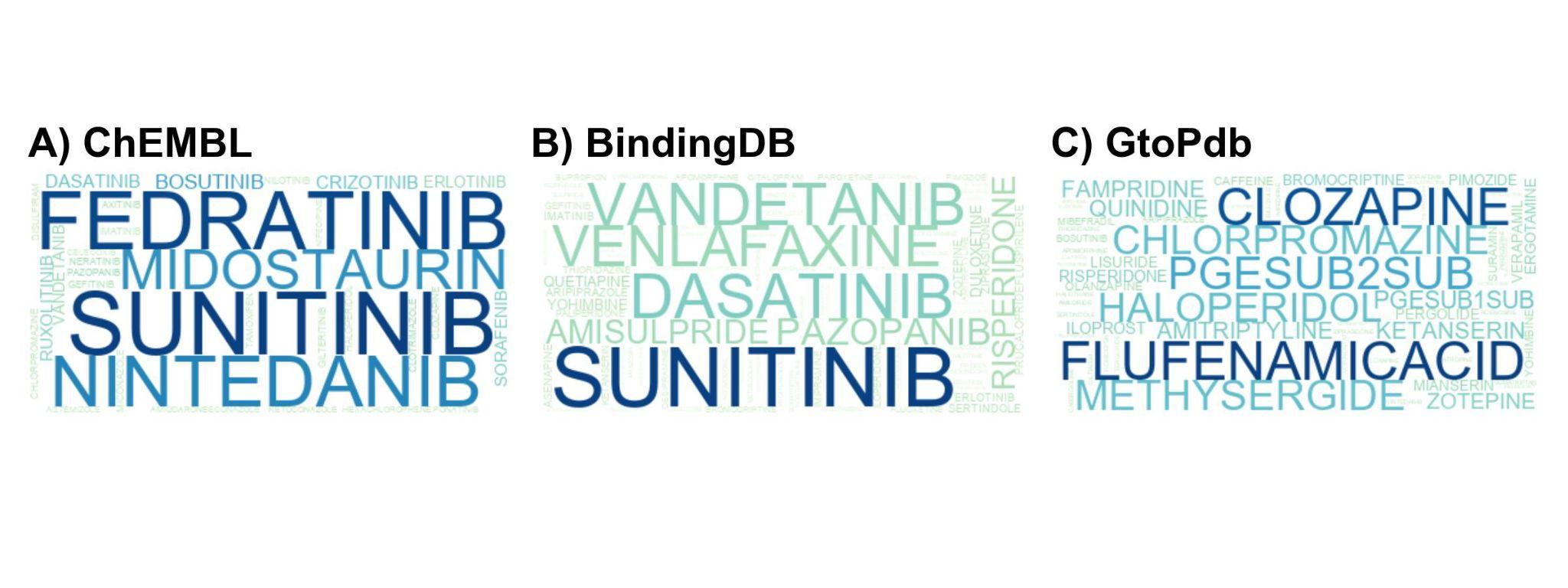

**Supplementary Figure 6:** Top 50 approved drugs from **A)** ChEMBL, **B)** BindingDB, and **C)** GtoPdb. The font size of each drug name reflects the number of targets associated with it in the respective database. An activity threshold of pChEMBL ≥0 was applied for inclusion in this analysis.

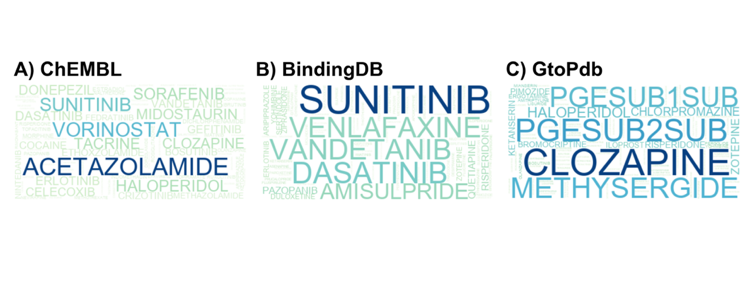

**Supplementary Figure 7:** Top 50 approved drugs from **A)** ChEMBL, **B)** BindingDB, and **C)** GtoPdb. The font size of each drug name reflects the number of targets associated with it in the respective database. An activity threshold of pChEMBL ≥5 was applied for inclusion in this analysis.

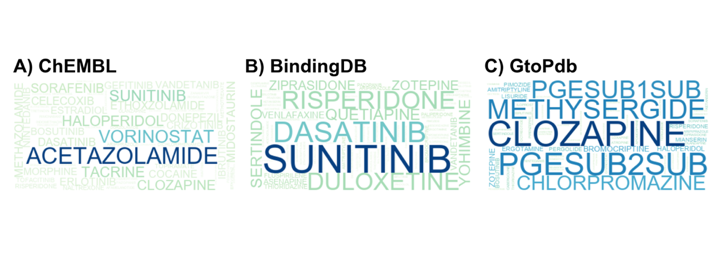

**Supplementary Figure 8:** Top 50 approved drugs from **A)** ChEMBL, **B)** BindingDB, and **C)** GtoPdb. The font size of each drug name reflects the number of targets associated with it in the respective database. An activity threshold of pChEMBL ≥6 was applied while doing this analysis.

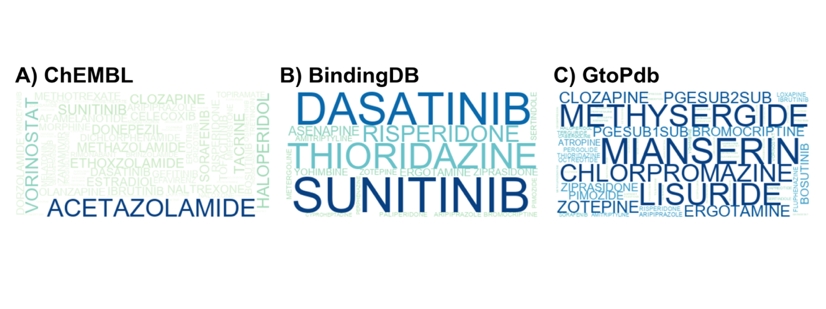

**Supplementary Figure 9:** Top 50 approved drugs from **A)** ChEMBL, **B)** BindingDB, and **C)** GtoPdb. The font size of each drug name reflects the number of targets associated with it in the respective database. An activity threshold of pChEMBL ≥7 was applied while doing this analysis.

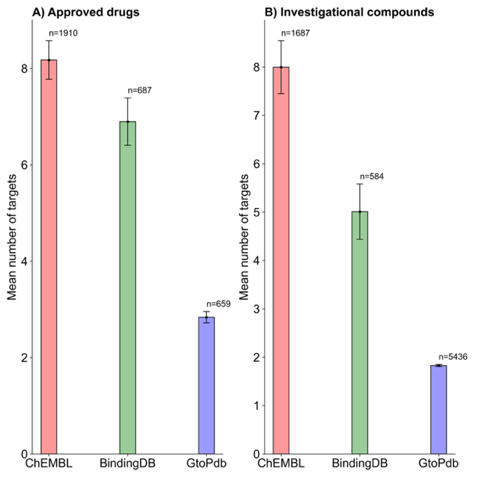

**Supplementary Figure 10**: Mean number of targets across the three databases at a pChEMBL ≥ 5. **(A)** Approved drugs; **(B)** Investigational compounds. “n” indicates the total number of drugs or compounds included in each analysis.

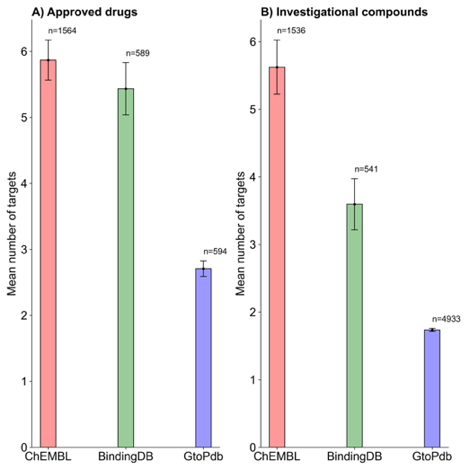

**Supplementary Figure 11** Mean number of targets across the three databases at a pChEMBL ≥ 6. **(A)** Approved drugs; **(B)** Investigational compounds. “n” indicates the total number of drugs or compounds included in each analysis.

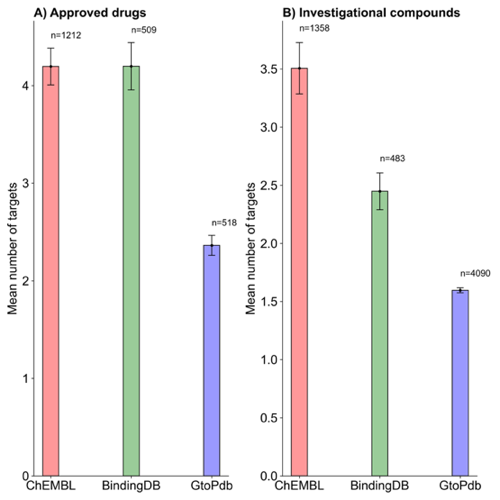

**Supplementary Figure 12**: Mean number of targets across the three databases at a pChEMBL ≥ 7. **(A)** Approved drugs; **(B)** Investigational compounds. “n” indicates the total number of drugs or compounds included in each analysis.

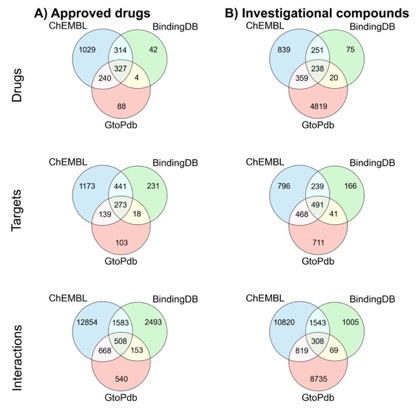

**Supplementary Figure 13**: Overlap of drugs, targets, and interactions across approved drugs and investigational compounds in the three binding affinity databases. Target overlap is determined using UniProt IDs, and compound overlap is based on standard InChIKey identifiers. A threshold of pChEMBL ≥5 is used for this analysis.

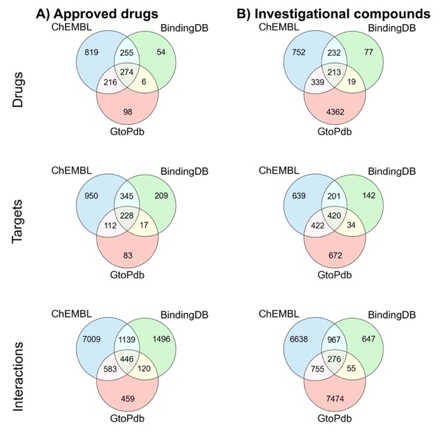

**Supplementary Figure 14**: Overlap of drugs, targets, and interactions across approved drugs and investigational compounds in the three binding affinity databases. Target overlap is determined using UniProt IDs, and compound overlap is based on standard InChIKey identifiers. A threshold of pChEMBL ≥6 is used for this analysis.

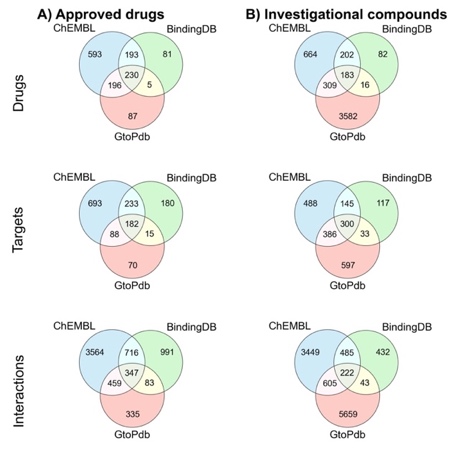

**Supplementary Figure 15**: Overlap of drugs, targets, and interactions across approved drugs and investigational compounds in the three binding affinity databases. Target overlap is determined using UniProt IDs, and compound overlap is based on standard InChIKey identifiers. A threshold of pChEMBL ≥7 is used for this analysis.

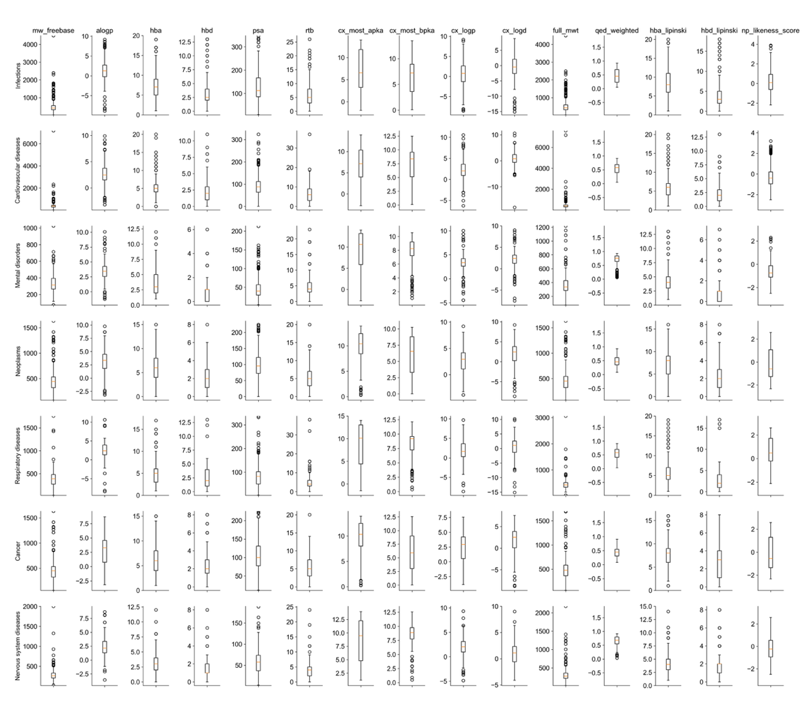

**Supplementary Figure 16**: Distribution of 15 physicochemical properties (columns) for approved drugs across 7 indication groups (rows): infections, cardiovascular, mental, neoplasms, respiratory, cancer, and nervous system diseases. Orange lines represent the median values for each indication group across the 15 properties.

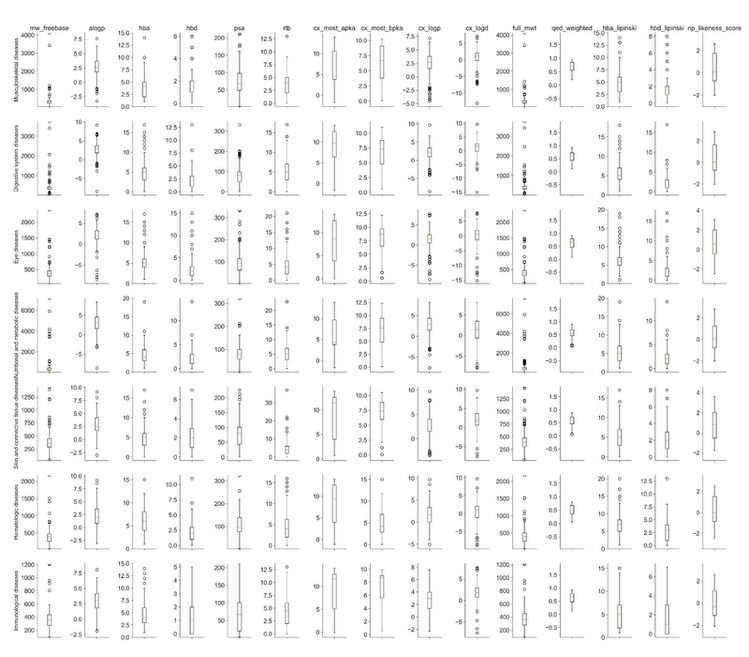

**Supplementary Figure 17:** Distribution of 15 physicochemical properties (columns) for approved drugs across indication groups: musculoskeletal, digestive system, eye, nutritional and metabolic, skin and connective tissue, hematologic and immunological. Orange lines represent the median values for each indication group across the 15 properties.

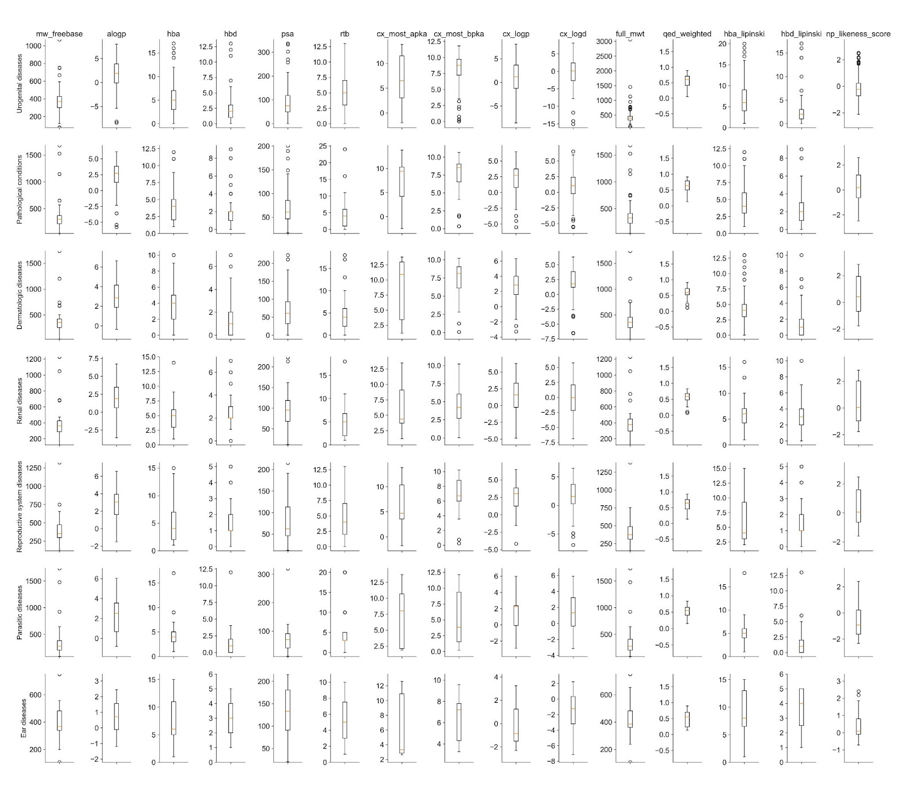

**Supplementary Figure 18:** Distribution of 15 physicochemical properties (columns) for approved drugs across indication groups: urogenital diseases, pathological conditions, dermatologic, renal, reproductive system, parasitic and ear.

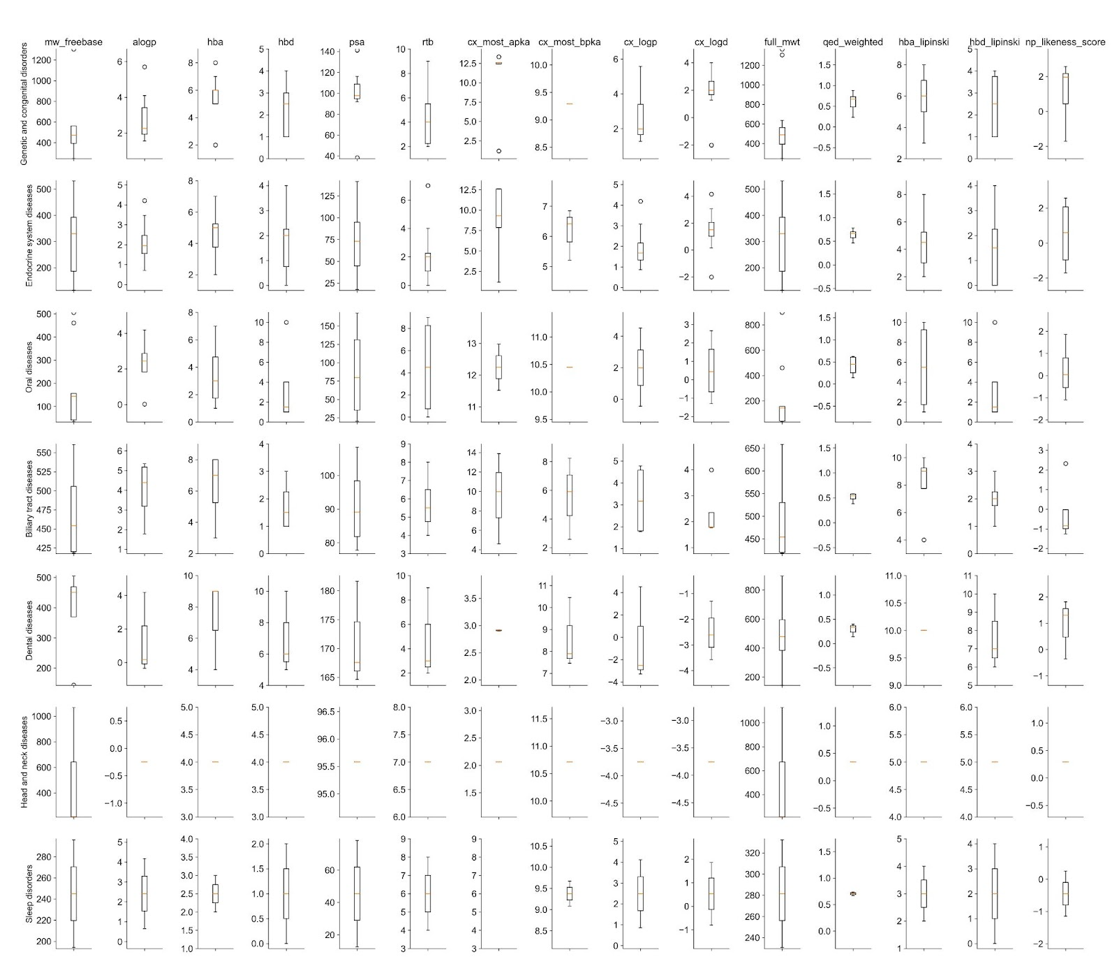

**Supplementary Figure 19:** Distribution of 15 physicochemical properties (columns) for approved drugs across indication groups: genetic and congenital, dental, endocrine system, head and neck, oral diseases, biliary tract and sleep disorders.

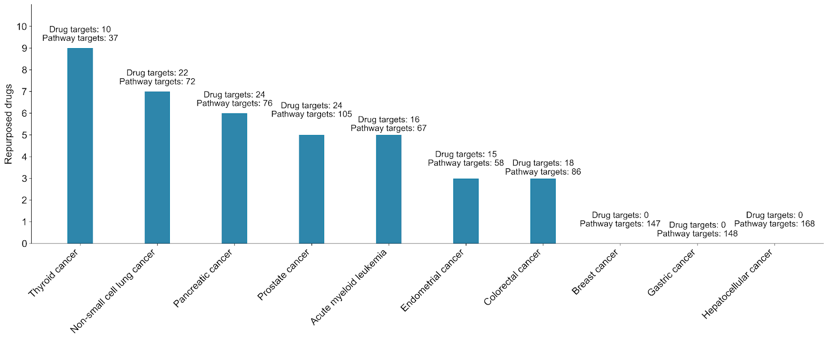

**Supplementary Figure 20:** Number of repurposed drugs identified across ten cancer types (at pChEMBL ≥5 with at least 10% pathway coverage). The labels above each bar indicate the total number of pathway-associated targets for each cancer type, as well as the subset of those targets that are known to bind repurposed drugs.

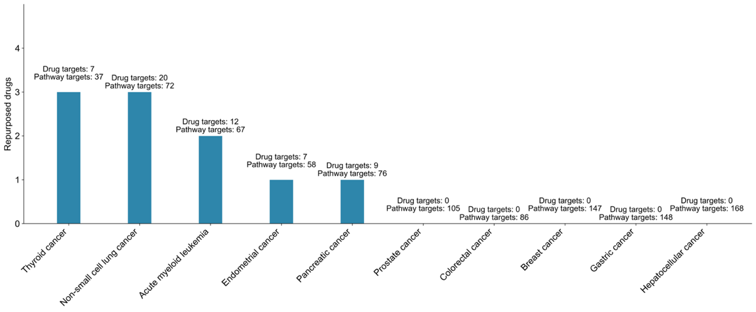

**Supplementary Figure 21:** Number of repurposed drugs identified across ten cancer types (at pChEMBL ≥6 with at least 10% pathway coverage). The labels above each bar indicate the total number of pathway-associated targets for each cancer type, as well as the subset of those targets that are known to bind repurposed drugs.

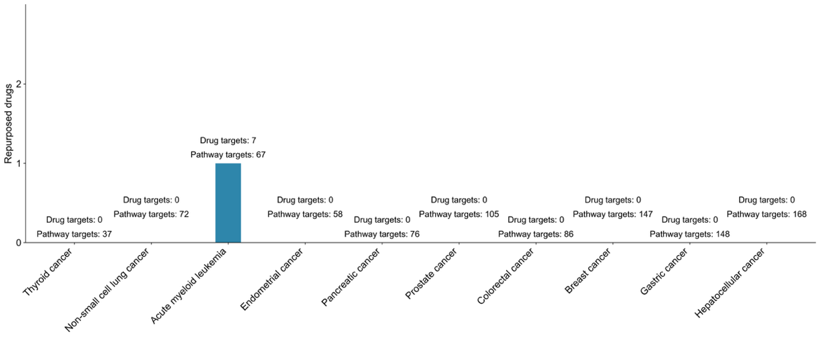

**Supplementary Figure 22:** Number of repurposed drugs identified across ten cancer types (at pChEMBL ≥7 with at least 10% pathway coverage). The labels above each bar indicate the total number of pathway-associated targets for each cancer type, as well as the subset of those targets that are known to bind repurposed drugs.

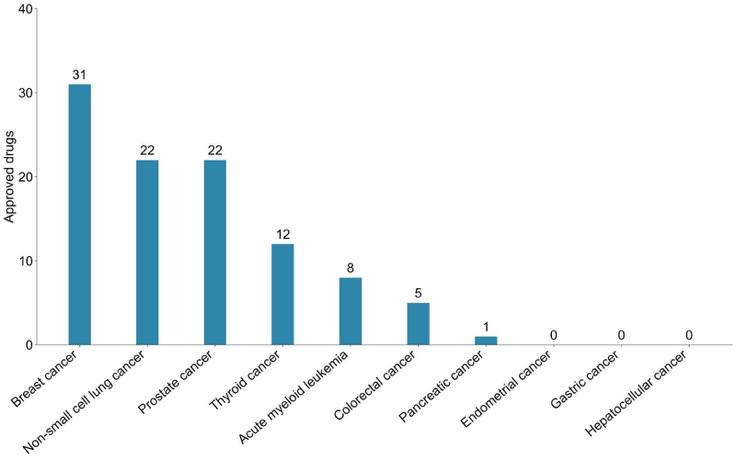

**Supplementary Figure 23:** Number of approved drugs across 10 cancer types as reported in ChEMBL (V33).

**Supplementary Table 1:** Targets involved in KEGG pathways for ten cancer types, with pathway IDs listed beneath each cancer type in the first column. Gene targets not associated with reviewed UniProt entries are removed.

| **Disease pathways** | **Target genes in the pathways** |
| --- | --- |
| Acute myeloid leukemia (hsa05221) | AKT1,AKT2,AKT3,ARAF,BAD,BCL2A1,BRAF,CCNA1,CCNA2,CCND1,CD14,CEBPA,CEBPE,CHUK,CSF1R,CSF2,DUSP6,EIF4EBP1,FCGR1A,FLT3,GRB2,HRAS,IKBKB,IKBKG,IL3,ITGAM,JUP,KIT,KRAS,LEF1,MAP2K1,MAP2K2,MAPK1,MAPK3,MPO,MTOR,MYC,NFKB1,NRAS,PER2,PIK3CA,PIK3CB,PIK3CD,PIK3R1,PIK3R2,PIK3R3,PIM1,PIM2,PML,PPARD,RAF1,RARA,RELA,RPS6KB1,RPS6KB2,RUNX1,RUNX1T1,SOS1,SOS2,SPI1,STAT3,STAT5A,STAT5B,TCF7,TCF7L1,TCF7L2,ZBTB16 |
| Breast cancer (hsa05224) | AKT1,AKT2,AKT3,APC,APC2,ARAF,AXIN1,AXIN2,BAK1,BAX,BRAF,BRCA1,BRCA2,CCND1,CDK4,CDK6,CDKN1A,CSNK1A1,CSNK1A1L,CTNNB1,DDB2,DLL1,DLL3,DLL4,DVL1,DVL2,DVL3,E2F1,E2F2,E2F3,EGF,EGFR,ERBB2,ESR1,ESR2,FGF1,FGF10,FGF16,FGF17,FGF18,FGF19,FGF2,FGF20,FGF21,FGF22,FGF23,FGF3,FGF4,FGF5,FGF6,FGF7,FGF8,FGF9,FGFR1,FLT4,FOS,FRAT1,FRAT2,FZD1,FZD10,FZD2,FZD3,FZD4,FZD5,FZD6,FZD7,FZD8,FZD9,GADD45A,GADD45B,GADD45G,GRB2,GSK3B,HES1,HES5,HEY1,HEY2,HEYL,HRAS,IGF1,IGF1R,JAG1,JAG2,JUN,KIT,KRAS,LEF1,LRP5,LRP6,MAP2K1,MAP2K2,MAPK1,MAPK3,MTOR,MYC,NCOA1,NCOA3,NFKB2,NOTCH1,NOTCH2,NOTCH3,NOTCH4,NRAS,PGR,PIK3CA,PIK3CB,PIK3CD,PIK3R1,PIK3R2,PIK3R3,POLK,PTEN,RAF1,RB1,RPS6KB1,RPS6KB2,SHC1,SHC2,SHC3,SHC4,SOS1,SOS2,SP1,TCF7,TCF7L1,TCF7L2,TNFSF11,TP53,WNT1,WNT10A,WNT10B,WNT11,WNT16,WNT2,WNT2B,WNT3,WNT3A,WNT4,WNT5A,WNT5B,WNT6,WNT7A,WNT7B,WNT8A,WNT8B,WNT9A,WNT9B |
| Colorectal cancer (hsa05210) | AKT1,AKT2,AKT3,APC,APC2,APPL1,ARAF,AREG,AXIN1,AXIN2,BAD,BAK1,BAX,BBC3,BCL2,BCL2L11,BIRC5,BRAF,CASP3,CASP9,CCND1,CDKN1A,CTNNB1,CYCS,DCC,DDB2,EGF,EGFR,EREG,FOS,GADD45A,GADD45B,GADD45G,GRB2,GSK3B,HRAS,JUN,KRAS,LEF1,MAP2K1,MAP2K2,MAPK1,MAPK10,MAPK3,MAPK8,MAPK9,MLH1,MSH2,MSH3,MSH6,MTOR,MYC,NRAS,PIK3CA,PIK3CB,PIK3CD,PIK3R1,PIK3R2,PIK3R3,PMAIP1,POLK,RAC1,RAC2,RAC3,RAF1,RALA,RALB,RALGDS,RHOA,RPS6KB1,RPS6KB2,SMAD2,SMAD3,SMAD4,SOS1,SOS2,TCF7,TCF7L1,TCF7L2,TGFA,TGFB1,TGFB2,TGFB3,TGFBR1,TGFBR2,TP53 |
| Endometrial cancer (hsa05213) | AKT1,AKT2,AKT3,APC,APC2,ARAF,AXIN1,AXIN2,BAD,BAK1,BAX,BRAF,CASP9,CCND1,CDH1,CDKN1A,CTNNA1,CTNNA2,CTNNA3,CTNNB1,DDB2,EGF,EGFR,ELK1,ERBB2,FOXO3,GADD45A,GADD45B,GADD45G,GRB2,GSK3B,HRAS,ILK,KRAS,LEF1,MAP2K1,MAP2K2,MAPK1,MAPK3,MLH1,MYC,NRAS,PDPK1,PIK3CA,PIK3CB,PIK3CD,PIK3R1,PIK3R2,PIK3R3,POLK,PTEN,RAF1,SOS1,SOS2,TCF7,TCF7L1,TCF7L2,TP53 |
| Gastric cancer (hsa05226) | ABCB1,AKT1,AKT2,AKT3,APC,APC2,ARAF,AXIN1,AXIN2,BAK1,BAX,BCL2,BRAF,CCND1,CCNE1,CCNE2,CDH1,CDH17,CDK2,CDKN1A,CDKN1B,CDKN2B,CDX2,CSNK1A1,CSNK1A1L,CTNNA1,CTNNA2,CTNNA3,CTNNB1,DDB2,DVL1,DVL2,DVL3,E2F1,E2F2,E2F3,EGF,EGFR,ERBB2,FGF1,FGF10,FGF16,FGF17,FGF18,FGF19,FGF2,FGF20,FGF21,FGF22,FGF23,FGF3,FGF4,FGF5,FGF6,FGF7,FGF8,FGF9,FGFR2,FRAT1,FRAT2,FZD1,FZD10,FZD2,FZD3,FZD4,FZD5,FZD6,FZD7,FZD8,FZD9,GAB1,GADD45A,GADD45B,GADD45G,GRB2,GSK3B,HGF,HRAS,JUP,KRAS,LEF1,LRP5,LRP6,MAP2K1,MAP2K2,MAPK1,MAPK3,MET,MLH1,MTOR,MUC2,MYC,NRAS,PIK3CA,PIK3CB,PIK3CD,PIK3R1,PIK3R2,PIK3R3,POLK,RAF1,RARB,RB1,REG4,RPS6KB1,RPS6KB2,RXRA,RXRB,RXRG,SHC1,SHC2,SHC3,SHC4,SHH,SMAD2,SMAD3,SMAD4,SOS1,SOS2,TCF7,TCF7L1,TCF7L2,TERT,TGFB1,TGFB2,TGFB3,TGFBR1,TGFBR2,TP53,WNT1,WNT10A,WNT10B,WNT11,WNT16,WNT2,WNT2B,WNT3,WNT3A,WNT4,WNT5A,WNT5B,WNT6,WNT7A,WNT7B,WNT8A,WNT8B,WNT9A,WNT9B |
| Hepatocellular carcinoma (hsa05225) | ACTB,ACTG1,ACTL6A,ACTL6B,AKT1,AKT2,AKT3,APC,APC2,ARAF,ARID1A,ARID1B,ARID2,AXIN1,AXIN2,BAD,BAK1,BAX,BCL2L1,BRAF,BRD7,CCND1,CDK4,CDK6,CDKN1A,CDKN2A,CSNK1A1,CSNK1A1L,CTNNB1,DDB2,DPF1,DPF3,DVL1,DVL2,DVL3,E2F1,E2F2,E2F3,EGFR,ELK1,FRAT1,FRAT2,FZD1,FZD10,FZD2,FZD3,FZD4,FZD5,FZD6,FZD7,FZD8,FZD9,GAB1,GADD45A,GADD45B,GADD45G,GRB2,GSK3B,GSTA1,GSTA2,GSTA3,GSTA4,GSTA5,GSTM1,GSTM2,GSTM3,GSTM4,GSTM5,GSTO1,GSTO2,GSTP1,GSTT1,GSTT2,GSTT2B,GSTT4,HGF,HMOX1,HRAS,IGF1R,IGF2,KEAP1,KRAS,LEF1,LRP5,LRP6,MAP2K1,MAP2K2,MAPK1,MAPK3,MET,MGST1,MGST2,MGST3,MTOR,MYC,NFE2L2,NQO1,NRAS,PBRM1,PHF10,PIK3CA,PIK3CB,PIK3CD,PIK3R1,PIK3R2,PIK3R3,PLCG1,PLCG2,POLK,PRKCA,PRKCB,PRKCG,PTEN,RAF1,RB1,RPS6KB1,RPS6KB2,SHC1,SHC2,SHC3,SHC4,SMAD2,SMAD3,SMAD4,SMARCA2,SMARCA4,SMARCB1,SMARCC1,SMARCC2,SMARCD1,SMARCD2,SMARCD3,SMARCE1,SOS1,SOS2,TCF7,TCF7L1,TCF7L2,TERT,TGFA,TGFB1,TGFB2,TGFB3,TGFBR1,TGFBR2,TP53,TXNRD1,TXNRD2,TXNRD3,WNT1,WNT10A,WNT10B,WNT11,WNT16,WNT2,WNT2B,WNT3,WNT3A,WNT4,WNT5A,WNT5B,WNT6,WNT7A,WNT7B,WNT8A,WNT8B,WNT9A,WNT9B |
| Non-small cell lung cancer (hsa05223) | AKT1,AKT2,AKT3,ALK,ARAF,BAD,BAK1,BAX,BRAF,CASP9,CCND1,CDK4,CDK6,CDKN1A,CDKN2A,DDB2,E2F1,E2F2,E2F3,EGF,EGFR,EML4,ERBB2,FHIT,FOXO3,GADD45A,GADD45B,GADD45G,GRB2,HGF,HRAS,JAK3,KIF5A,KIF5B,KIF5C,KRAS,MAP2K1,MAP2K2,MAPK1,MAPK3,MET,NRAS,PDPK1,PIK3CA,PIK3CB,PIK3CD,PIK3R1,PIK3R2,PIK3R3,PLCG1,PLCG2,POLK,PRKCA,PRKCB,PRKCG,RAF1,RARB,RASSF1,RASSF5,RB1,RET,RXRA,RXRB,RXRG,SOS1,SOS2,STAT3,STAT5A,STAT5B,STK4,TGFA,TP53 |
| Pancreatic cancer (hsa05212) | AKT1,AKT2,AKT3,ARAF,ARHGEF6,BAD,BAK1,BAX,BCL2L1,BRAF,BRCA2,CASP9,CCND1,CDC42,CDK4,CDK6,CDKN1A,CDKN2A,CHUK,DDB2,E2F1,E2F2,E2F3,EGF,EGFR,ERBB2,GADD45A,GADD45B,GADD45G,IKBKB,IKBKG,JAK1,KRAS,MAP2K1,MAPK1,MAPK10,MAPK3,MAPK8,MAPK9,MTOR,NFKB1,PIK3CA,PIK3CB,PIK3CD,PIK3R1,PIK3R2,PIK3R3,PLD1,PLD2,POLK,RAC1,RAC2,RAC3,RAD51,RAF1,RALA,RALB,RALBP1,RALGDS,RB1,RELA,RPS6KB1,RPS6KB2,SMAD2,SMAD3,SMAD4,STAT1,STAT3,TGFA,TGFB1,TGFB2,TGFB3,TGFBR1,TGFBR2,TP53,VEGFA |
| Prostate cancer (hsa05215) | AKT1,AKT2,AKT3,AR,ARAF,ATF4,BAD,BCL2,BRAF,CASP9,CCND1,CCNE1,CCNE2,CDK2,CDKN1A,CDKN1B,CHUK,CREB1,CREB3,CREB3L1,CREB3L2,CREB3L3,CREB3L4,CREB5,CREBBP,CTNNB1,E2F1,E2F2,E2F3,EGF,EGFR,EP300,ERBB2,ERG,ETV5,FGFR1,FGFR2,FOLH1,FOLR1,FOLR2,FOLR3,FOXO1,GRB2,GRM1,GRM5,GSK3B,GSTP1,HRAS,HSP90AA1,HSP90AB1,HSP90B1,IGF1,IGF1R,IKBKB,IKBKG,IL1R2,INS,INSRR,KLK3,KRAS,LEF1,MAP2K1,MAP2K2,MAPK1,MAPK3,MDM2,MMP3,MMP9,MTOR,NFKB1,NFKBIA,NKX3-1,NRAS,PDGFA,PDGFB,PDGFC,PDGFD,PDGFRA,PDGFRB,PDPK1,PIK3CA,PIK3CB,PIK3CD,PIK3R1,PIK3R2,PIK3R3,PLAT,PLAU,PTEN,RAF1,RB1,RELA,SLC19A1,SLC46A1,SOS1,SOS2,SPINT1,SRD5A2,TCF7,TCF7L1,TCF7L2,TGFA,TMPRSS2,TP53,ZEB1 |
| Thyroid cancer (hsa05216) | BAK1,BAX,BRAF,CCDC6,CCND1,CDH1,CDKN1A,CTNNB1,DDB2,GADD45A,GADD45B,GADD45G,HRAS,KRAS,LEF1,MAP2K1,MAP2K2,MAPK1,MAPK3,MYC,NCOA4,NRAS,NTRK1,PAX8,POLK,PPARG,RET,RXRA,RXRB,RXRG,TCF7,TCF7L1,TCF7L2,TFG,TP53,TPM3,TPR |

**Supplementary Table 2:** Repurposed drugs for different cancer types identified using a potency threshold of pChEMBL ≥ 6 and requiring that drug targets overlap with at least 10% of the targets in the corresponding cancer-related pathways.

| **Cancer** | **Repurposing drug** | **Targets in pathway** | **Targets binding to drug** | **Binding targets** |
| --- | --- | --- | --- | --- |
| Thyroid cancer | Nintedanib | 37 | 4 | MAP2K1, NTRK1, RET, MAP2K2 |
| Thyroid cancer | Sorafenib | 37 | 4 | MAPK1, MAPK3, RET, BRAF |
| Thyroid cancer | Sunitinib | 37 | 4 | MAP2K1, NTRK1, RET, MAP2K2 |
| Non-small cell lung cancer | Midostaurin | 72 | 9 | JAK3, PRKCA, AKT1, ALK, PDPK1, STK4, AKT2, MET, RET |
| Non-small cell lung cancer | Nintedanib | 72 | 8 | MAP2K1, JAK3, ALK, MAP2K2, STK4, MET, RET, CDK4 |
| Non-small cell lung cancer | Sorafenib | 72 | 9 | EGFR, CDK6, MAPK3, MAPK1, RET, ARAF, BRAF, ERBB2, RAF1 |
| Acute myeloid leukemia | Sorafenib | 67 | 9 | RPS6KB1, MAPK3, KIT, CSF1R, MAPK1, ARAF, BRAF, FLT3, RAF1 |
| Acute myeloid leukemia | Sunitinib | 67 | 7 | RPS6KB1, CHUK, MAP2K1, KIT, CSF1R, MAP2K2, FLT3 |
| Endometrial cancer | Sorafenib | 58 | 7 | EGFR, MAPK3, MAPK1, ARAF, BRAF, ERBB2, RAF1 |
| Pancreatic cancer | Sorafenib | 76 | 9 | RPS6KB1, EGFR, CDK6, MAPK3, MAPK1, ARAF, BRAF, ERBB2, RAF1 |

**Supplementary Table 3:** Repurposed drugs for different cancer types identified using a potency threshold of pChEMBL ≥ 5 and requiring that drug targets overlap with at least 10% of the targets in the corresponding cancer-related pathways.

| **Cancer** | **Repurposing drug** | **Targets in pathway** | **Targets binding to drug** | **Binding targets** |
| --- | --- | --- | --- | --- |
| Prostate cancer | Fedratinib | 105 | 13 | PIK3CA, INSRR, IGF1R, EGFR, FGFR2, PDGFRB, BRAF, PDGFRA, MAP2K2, CHUK, IKBKB, FGFR1, MAP2K1 |
| Prostate cancer | Midostaurin | 105 | 13 | PIK3CA, AKT1, AKT3, PDGFRB, FGFR2, PDGFRA, GSK3B, CHUK, FGFR1, IKBKB, AKT2, EGFR, PDPK1 |
| Prostate cancer | Nintedanib | 105 | 12 | INSRR, CDK2, IGF1R, FGFR2, PDGFRB, PDGFRA, GSK3B, MAP2K2, CHUK, FGFR1, MAP2K1, PDPK1 |
| Prostate cancer | Sorafenib | 105 | 12 | MAPK1, ERBB2, ARAF, CDK2, FGFR2, PDGFRB, BRAF, PDGFRA, FGFR1, MAPK3, EGFR, RAF1 |
| Prostate cancer | Sunitinib | 105 | 12 | INSRR, IGF1R, EGFR, FGFR2, PDGFRB, PDGFRA, MAP2K2, CHUK, FGFR1, AKT2, MAP2K1, PDPK1 |
| Endometrial cancer | Dasatinib | 58 | 6 | ERBB2, EGFR, BRAF, MAP2K2, MAP2K1, RAF1 |
| Endometrial cancer | Midostaurin | 58 | 7 | PIK3CA, AKT1, AKT3, GSK3B, AKT2, EGFR, PDPK1 |
| Endometrial cancer | Sorafenib | 58 | 7 | MAPK1, ERBB2, ARAF, BRAF, MAPK3, EGFR, RAF1 |
| Non-small cell lung cancer | Bosutinib | 72 | 9 | ERBB2, MET, JAK3, RET, STK4, ALK, MAP2K2, MAP2K1, EGFR |
| Non-small cell lung cancer | Dasatinib | 72 | 9 | ERBB2, JAK3, BRAF, RET, STK4, MAP2K2, MAP2K1, EGFR, RAF1 |
| Non-small cell lung cancer | Fedratinib | 72 | 10 | MET, PIK3CA, JAK3, BRAF, RET, ALK, MAP2K2, CDK4, MAP2K1, EGFR |
| Non-small cell lung cancer | Midostaurin | 72 | 12 | PRKCA, MET, PIK3CA, AKT1, AKT3, JAK3, RET, STK4, ALK, AKT2, EGFR, PDPK1 |
| Non-small cell lung cancer | Nintedanib | 72 | 9 | MET, JAK3, RET, STK4, ALK, MAP2K2, CDK4, MAP2K1, PDPK1 |
| Non-small cell lung cancer | Sorafenib | 72 | 11 | MAPK1, ERBB2, MET, ARAF, JAK3, BRAF, RET, CDK6, MAPK3, EGFR, RAF1 |
| Non-small cell lung cancer | Sunitinib | 72 | 12 | PRKCA, MET, MAP2K1, JAK3, RET, STK4, ALK, MAP2K2, CDK4, AKT2, EGFR, PDPK1 |
| Colorectal cancer | Fedratinib | 86 | 10 | PIK3CA, RPS6KB1, EGFR, TGFBR2, BRAF, MAP2K2, MAPK8, MAPK10, MAPK9, MAP2K1 |
| Colorectal cancer | Midostaurin | 86 | 10 | PIK3CA, RPS6KB1, AKT1, AKT3, GSK3B, MAPK10, MAPK8, MAPK9, AKT2, EGFR |
| Colorectal cancer | Sorafenib | 86 | 9 | MAPK1, ARAF, RPS6KB1, TGFBR2, BRAF, MAPK9, MAPK3, EGFR, RAF1 |
| Pancreatic cancer | Dabrafenib | 76 | 8 | ARAF, BRAF, TGFBR2, CDK6, CDK4, TGFBR1, RAF1, KRAS |
| Pancreatic cancer | Fedratinib | 76 | 13 | PIK3CA, RPS6KB1, BRAF, TGFBR2, CHUK, CDK4, MAPK8, MAPK9, IKBKB, MAP2K1, EGFR, MAPK10, JAK1 |
| Pancreatic cancer | Midostaurin | 76 | 12 | PIK3CA, RPS6KB1, AKT1, AKT3, CHUK, MAPK8, IKBKB, MAPK9, JAK1, AKT2, EGFR, MAPK10 |
| Pancreatic cancer | Nintedanib | 76 | 8 | RPS6KB1, CDK4, CHUK, MAPK8, TGFBR1, JAK1, MAPK10, MAP2K1 |
| Pancreatic cancer | Sorafenib | 76 | 11 | MAPK1, ERBB2, ARAF, RPS6KB1, BRAF, TGFBR2, CDK6, MAPK9, MAPK3, EGFR, RAF1 |
| Pancreatic cancer | Sunitinib | 76 | 9 | RPS6KB1, MAP2K1, CDK4, CHUK, MAPK10, MAPK9, JAK1, AKT2, EGFR |
| Thyroid cancer | Axitinib | 37 | 4 | MAP2K1, MAP2K2, NTRK1, RET |
| Thyroid cancer | Bosutinib | 37 | 4 | MAP2K1, MAP2K2, NTRK1, RET |
| Thyroid cancer | Dasatinib | 37 | 4 | MAP2K1, MAP2K2, BRAF, RET |
| Thyroid cancer | Fedratinib | 37 | 5 | BRAF, RET, MAP2K2, MAP2K1, NTRK1 |
| Thyroid cancer | Nintedanib | 37 | 4 | MAP2K1, MAP2K2, NTRK1, RET |
| Thyroid cancer | Ruxolitinib | 37 | 5 | BRAF, RET, MAP2K2, MAP2K1, NTRK1 |
| Thyroid cancer | Sorafenib | 37 | 5 | MAPK1, BRAF, RET, MAPK3, NTRK1 |
| Thyroid cancer | Sunitinib | 37 | 4 | MAP2K1, MAP2K2, NTRK1, RET |
| Thyroid cancer | Tretinoin | 37 | 4 | RXRG, MAPK1, RXRA, RXRB |
| Acute myeloid leukemia | Dasatinib | 67 | 7 | FLT3, BRAF, CSF1R, MAP2K2, MAP2K1, RAF1, KIT |
| Acute myeloid leukemia | Fedratinib | 67 | 10 | FLT3, PIK3CA, RPS6KB1, BRAF, CSF1R, MAP2K2, CHUK, IKBKB, MAP2K1, KIT |
| Acute myeloid leukemia | Nintedanib | 67 | 7 | FLT3, RPS6KB1, CSF1R, CHUK, MAP2K2, MAP2K1, KIT |
| Acute myeloid leukemia | Sorafenib | 67 | 9 | FLT3, MAPK1, ARAF, RPS6KB1, BRAF, CSF1R, MAPK3, RAF1, KIT |
| Acute myeloid leukemia | Sunitinib | 67 | 9 | FLT3, PIM2, RPS6KB1, CSF1R, MAP2K2, CHUK, AKT2, MAP2K1, KIT |

**Supplementary Table 4:** Approved drugs for ten cancer types as reported in ChEMBL (V 33).

| **Cancer** | **Approved drugs** |
| --- | --- |
| Breast cancer | Abemaciclib, alpelisib, anastrozole, capecitabine, cyclophosphamide, dexrazoxane hydrochloride, docetaxel, doxorubicin hydrochloride, epirubicin hydrochloride, eribulin mesylate, estradiol cypionate, everolimus, exemestane, fluoroestradiol f-18, fulvestrant, gemcitabine hydrochloride, goserelin acetate, ixabepilone, lapatinib ditosylate, letrozole, methotrexate, methyltestosterone, neratinib maleate, paclitaxel, palbociclib, raloxifene hydrochloride, ribociclib succinate, talazoparib tosylate, tamoxifen citrate, toremifene citrate, tucatinib |
| Non-small cell lung cancer | Afatinib dimaleate, alectinib hydrochloride, brigatinib, capmatinib hydrochloride, ceritinib, crizotinib, dacomitinib, docetaxel, entrectinib, erlotinib hydrochloride, gefitinib, gemcitabine hydrochloride, lorlatinib, mobocertinib succinate, osimertinib, osimertinib mesylate, paclitaxel, pemetrexed disodium, pralsetinib, selpercatinib, sotorasib, tepotinib hydrochloride |
| Pancreatic cancer | Gemcitabine hydrochloride |
| Prostate cancer | Abiraterone acetate, apalutamide, cabazitaxel, choline c-11, darolutamide, degarelix acetate, dutasteride, enzalutamide, finasteride, flutamide, histrelin acetate, isopropyl alcohol, leuprolide acetate, leuprolide mesylate, lidocaine hydrochloride, mitoxantrone hydrochloride, nilutamide, radium ra 223 dichloride, relugolix, tamsulosin hydrochloride, triptorelin pamoate, zoledronic acid anhydrous |
| Thyroid cancer | Cabozantinib s-malate, dabrafenib mesylate, lenvatinib, lenvatinib mesylate, levothyroxine sodium, liothyronine sodium, selpercatinib, sodium iodide i 123, sodium iodide i 131, sorafenib tosylate, trametinib dimethyl sulfoxide, vandetanib |
| Colorectal cancer | Levoleucovorin, levoleucovorin calcium, regorafenib, tipiracil hydrochloride, trifluridine |
| Acute Myeloid Leukemia | Cytarabine, daunorubicin, enasidenib mesylate, gilteritinib fumarate, glasdegib maleate, idarubicin hydrochloride, ivosidenib, midostaurin |
| Endometrial cancer | Nil |
| Gastric cancer | Nil |
| Hepatocellular cancer | Nil |

**Drug likeliness and Lipinski’s rule of 5 violations for drugs in ChEMBL**

The physicochemical properties of compounds are known to play a critical role in drug discovery and development, with certain parameters often considered essential for regulatory approval. Aromatic ring count is regarded as an important factor due to its influence on ligand solubility, which in turn affects target binding efficiency and elimination from the body. Depending on the study, the optimal number of aromatic rings within a drug molecule is recommended to be $\leq3$ or $\leq$ 5.

The calculated partition coefficient (LogP) serves as a key descriptor of lipophilicity, reflecting a compound’s affinity for nonpolar environments. Lipophilicity is recognized as a determinant of bioavailability, membrane permeability, and binding affinity. LogP is defined as the logarithmic ratio of a compound’s solubility in n-octanol to its solubility in water-saturated n-octanol. Elevated lipophilicity is generally associated with improved membrane permeability, although excessive values may increase toxicity risks.

Hydrogen bond donors (HBDs), typically defined as hydroxyl (OH) and amine (NH) groups, and hydrogen bond acceptors (HBAs), generally nitrogen (N) and oxygen (O) atoms, are considered relevant to solubility, permeability, and blood–brain barrier penetration. The polar surface area (PSA) of a compound is correlated with both drug absorption and target affinity. Molecular weight (MW) is also recognized as an important determinant, with different administration routes imposing distinct size constraints.

The acid dissociation constant (pKa) is defined as the negative base-10 logarithm of the acid-base dissociation constant (Ka) $pK_{a}=-log_{10}{(K}_{a}).$ Here, $K_{a}$ is defined by the concentrations of the ligand (L), the protein (P) and the ligand-protein complex (LP) as $K_{a}=\frac{[L][P]}{[LP]}$.

A widely cited guideline in small-molecule drug discovery is Lipinski’s Rule of Five (Ro5), introduced in 1997, which links favorable oral bioavailability to four criteria: ≤5 HBDs, ≤10 HBAs, MW < 500 Da, and LogP ≤ 5. Compounds violating more than one criterion are generally considered less likely to exhibit optimal absorption or permeability, although notable therapeutic exceptions exist.

Ro5 compliance was assessed using physicochemical data from ChEMBL. **Supplementary Figures 24 and 25** show distributions of the four Ro5 properties for approved and investigational compounds, with violations highlighted in red and compliant compounds in blue. We further quantified the number of compounds violating one, two, three, or all four thresholds (**Supplementary Figure 26**). Most approved and investigational drugs met the Ro5 criteria; however, 21.8% (761/3,492) of approved and 32.3% (1,168/3,618) of investigational compounds violated at least one property. These results support the utility of Ro5 while highlighting that compounds outside its boundaries may still hold therapeutic potential.

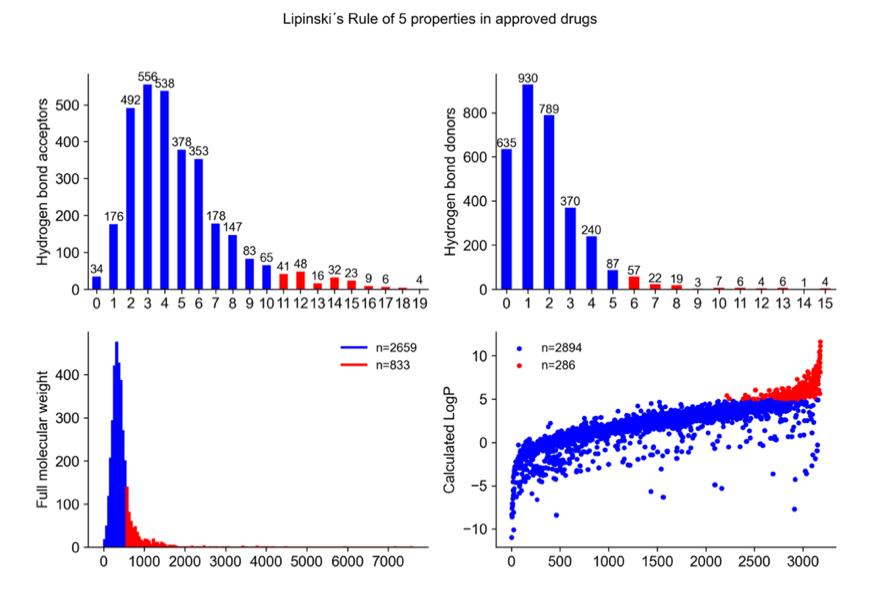

**Supplementary Figure 24:** Rule-of-five properties of approved drugs in the ChEMBL database. The blue color indicates compliance with the Ro5 threshold, while red color indicates violations. In the two upper graphs, the x-axes represents the number of hydrogen bond acceptors (left) or donors (right), with the number above the bar representing the number of approved drugs with that specific number of donors or acceptors. In the lower left graph, the x-axis value represents the molecular weight and the y-axis the frequency of drugs within that weight range. In the lower right graph, the y-axis represents the calculated LogP values, and the x-axis indicates the number of compounds corresponding to each value. The total number of Ro5-compliant and non-compliant drugs is shown in the top corner of each lower graph.

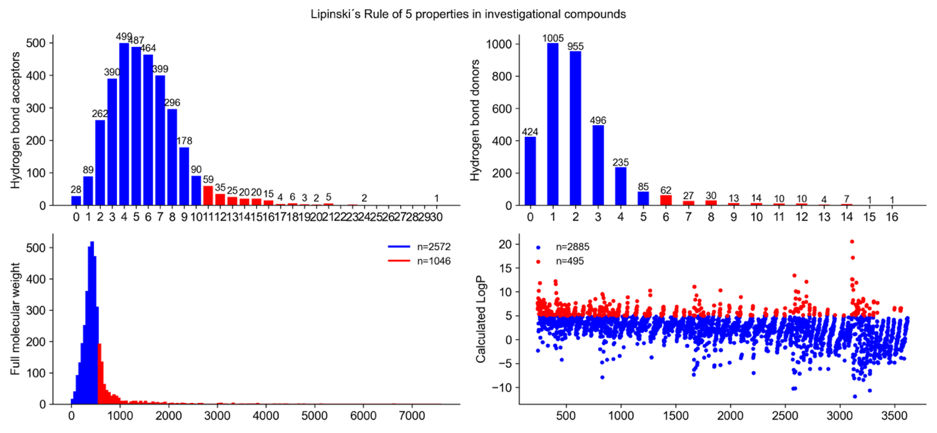

**Supplementary Figure 25:** Rule-of-five properties of investigational compounds in the ChEMBL database. The blue color indicates compliance with the Ro5 threshold, while red color indicates violations. In the two upper graphs, the x-axes represent the number of hydrogen bond acceptors (left) or donors (right), with the number above the bar representing the number of approved drugs with that specific number of donors or acceptors. In the lower left graph, the x-axis value represents the molecular weight and the y-axis the frequency of drugs within that weight range. In the lower right graph, the y-axis represents the calculated LogP values, and the x-axis indicates the number of compounds corresponding to each value. The total number of Ro5-compliant and non-compliant drugs is shown in the top corner of each lower graph.

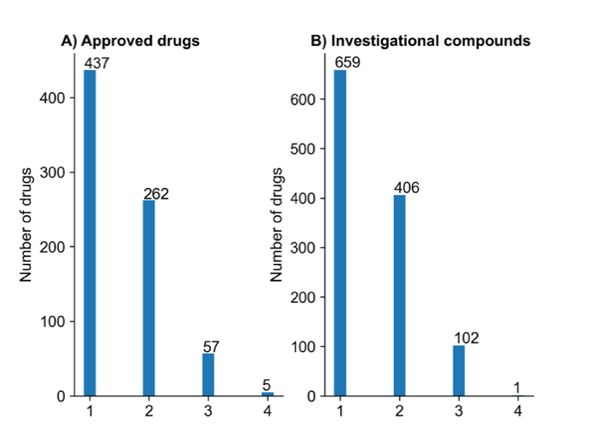

**Supplementary Figure 26:** Number of Lipinski’s Rule-of-Five (Ro5) violations among approved drugs (left) and investigational compounds (right). The x-axis indicates the number of Ro5 properties violated. The values above each bar represent the total number of compounds with that specific number of violations. The total number of approved drugs and investigational compounds is shown above each respective graph.
